# E prostanoid receptor 4 expressing macrophages promote the regeneration of the intestinal epithelial barrier upon inflammation

**DOI:** 10.1101/2020.05.04.077412

**Authors:** Yi Rang Na, Daun Jung, Michelle Stakenborg, Gyo Jeong Gu, Mi Reu Jeong, Hye Ri Jang, Soo Youn Suh, Hak Jae Kim, Yoon Hey Kwon, Tae Sik Sung, Seung Bum Ryoo, Kyu Joo Park, Jong Pil Im, Ji Yong Park, Yun Sang Lee, Heon Jong Han, Bo Youn Park, Sung Wook Lee, Ho Su Lee, Isabelle Cleynen, Gianluca Matteoli, Seung Hyeok Seok

## Abstract

Dysfunctional resolution of intestinal inflammation and altered mucosal healing are essential features in the pathogenesis of inflammatory bowel disease (IBD). Intestinal macrophages are vital in the process of resolution of inflammation but the mechanisms underlying their mucosal healing capacity remains elusive. Here, we describe a subset of E prostanoid receptor 4 (EP4) expressing intestinal macrophages with mucosal healing properties both in human and mice. Notably, Csf1r-iCre EP4-fl/fl mice showed defective mucosal healing and intestinal epithelial barrier regeneration in a dextran sodium sulfate-induced colitis model. Mechanistically, an increased mucosal level of prostaglandin E2 (PGE2) triggers the secretion of chemokine (C-X-C motif) ligand 1 (CXCL1) in monocyte-derived EP4^+^ macrophages via MAPKs. Subsequently, CXCL1 drives epithelial cell differentiation and proliferation from regenerating crypts during the resolution phase of colitis. Thus, EP4^+^ intestinal macrophages are essential for the support of the intestinal stem cell niche and for the regeneration of the injured epithelium.

**One Sentence Summary:** Prostaglandin E2 licenses E-type prostanoid receptor 4 intestinal macrophage regenerative capacity promoting mucosal healing via the secretion of CXCL1

## Main Text

Intestinal mucosal healing is the ultimate clinical criteria for complete remission in patients with inflammatory bowel disease (IBD) (*1*). The anatomical uniqueness of the intestine makes mucosal healing one of the most important requisites of therapeutic efficacy in IBD because of its role in preventing direct contact between the immune system, the microbiota and food antigens (*2*). In homeostasis, the epithelial barrier is maintained by highly active stem cells residing at the base of the crypts that provide continuous regeneration of enterocytes, with the majority of differentiated cells being replaced every 3-5 days (*3*). In response to injury, additional protective mechanisms are rapidly activated to restore the epithelial barrier (*4*). Delay or failure of intestinal epithelial restoration can facilitate exposure to luminal antigens and potentially drive invasion of luminal microorganisms into the mucosa, resulting in a proinflammatory response, tissue damage, and systemic infection. However, current therapeutic strategies are primarily aimed at altering the immune responses and do not directly promote mucosal healing.

Among the many anti-inflammatory drugs, nonsteroidal anti-inflammatory drugs (NSAIDs) are contraindicated in gastrointestinal inflammatory disorders, as they block cyclooxygenase-1 and -2 (COX-1/2), which catalyze the synthesis of prostaglandins, essential mediators for the homeostasis of the enterocytes (*5*). Likewise, glucocorticoids, which reduce the synthesis of prostaglandins by suppressing phospholipase A_2_, COX-2, and microsomal prostaglandin E synthase 2 (mPGES_2_), have been reported to adversely affect the regeneration of the intestinal epithelium (*6–8*). Even if the role of prostaglandins as essential mediators in intestinal inflammation is well-defined (*9*), there are only few reports on their cellular targets and mechanisms of action. Delineating the detailed cellular and molecular mechanisms that modulate the healing process via prostaglandins might provide an opportunity to identify detailed components that normally regulate the proliferative activities of epithelial progenitors. Identifying these mechanisms would enable the development of new therapeutic approaches to favor remission in patients suffering from IBD.

In addition, the presence of macrophages is essential to re-establish the epithelial barrier (*10, 11*). The macrophage mannose receptor (MMR, CD206)-expressing macrophages, which are decreased in patients with IBD (*12–14*), have been reported to have mucosal healing properties (*15, 16*). However, the mechanism underlying the formation of CD206^+^ intestinal macrophages has yet to be elucidated.

Prostaglandin E_2_ (PGE_2_) is one of the downstream lipid mediators of COXs (*17*) associated with resolution of inflammation and tissue regeneration in the skin, muscle and gut. In the intestine, PGE_2_ induces the differentiation of epithelial stem cells to wound-associated epithelial cells through EP4 (*2, 4*). In the skin, the effect of PGE_2_ was recently reported to promote the polarization of alternatively activated macrophages and enhanced wound healing in a murine cutaneous wound healing model (*18*). However, the specific mode of action of PGE_2_ on wound healing macrophages in the intestine remains unclear. In our previous study, endogenous PGE_2_ was found to potentiate the anti-inflammatory phenotype of resident macrophages (*19*). In addition, inhibition of 15-hydroxyprostaglandin dehydrogenase (15-PGDH), a PG-degrading enzyme, stimulated tissue regeneration in dextran sodium sulfate (DSS) colitis (*20*). Based on these studies, we hypothesized that PGE_2_ may promote the wound healing capacity of intestinal macrophages leading to resolution of intestinal inflammation. Here, we found that specific deletion of the prostaglandin PGE_2_ receptor, EP4, in macrophages resulted in insufficient epithelial cell regeneration after DSS-induced colitis. Mechanistically, increased level of PGE_2_ induced secretion of C-X-C motif chemokine ligand 1 (CXCL1) by EP4^+^ macrophages via MAPK pathways leading to intestinal crypt regeneration. Finally, specific therapeutic targeting of macrophages with liposomes loaded with a MAPK agonist augmented the production of CXCL1 *in vivo* in macrophage-specific EP4-deficient mice, restoring their defective epithelial regeneration and favoring mucosal healing. Overall, we offer novel insights into the role of EP4 in inducing tissue protective macrophages in intestinal inflammation, which paves the way for the development of a new class of targets to treat and/or favor remission in patients with IBD.

## Results

### E prostanoid receptor 4 is specifically expressed in CD206^+^ intestinal macrophages

The 5p13.1 locus is a well-defined genetic susceptibility locus for Crohn’s disease (CD),and has been shown to modulate the expression of the *PTGER4* gene. *PTGER4* is the gene located closest to the susceptibility region, and codes for EP4 (*21*). By analyzing the public GEO database, we found that patients with CD, as well as patients with ulcerative colitis (UC), showed reduced expression levels of the *PTGER4* gene in colonic mucosa biopsies compared with healthy individuals (Fig. S1A). In contrast, colonic biopsies of patients with UC in remission re-gained *PTGER4* gene expression (Fig. S1A).

As EP4 signaling seems to be implicated in IBD, we aimed to define the cellular localization of EP4 in the intestine using confocal microcopy and flow cytometry in murine and human colonic biopsies. In the healthy murine colonic mucosa, we observed that the majority of EP4^+^ cells displayed a macrophage-like morphology (Fig. 1A) and were scattered throughout the lamina propria and near the epithelial crypts. In addition, immunofluorescent (IF) staining of EP4 in the colon of tamoxifen-inducible *Csf1r-creERT*;Ai14(*RCL-tdT*) mice revealed colocalization of EP4 with CSF1R^+^ tdT^+^ cells. To better understand the expression pattern of EP4 in myeloid cells, we analyzed the expression of EP4 via flow cytometry during differentiation of lamina propria monocytes (P1) into intermediate (P2) and mature macrophages (P3) in the healthy mouse colon (Fig. 1B). EP4 was not detected in monocytes, whereas half of the intermediate macrophages and around 70 % of mature macrophages expressed EP4, indicating that macrophages acquires EP4 expression during their differentiation. In accordance with the IF data, mature macrophages expressed the highest levels of EP4 among other mucosal immune cells including CD45^neg^ non-immune cells and CD45^+^ immune cells such as lymphocytes, neutrophils, dendritic cells, and monocytes (Fig. 1C; gating strategy is shown in Fig. S2).

**Fig. 1.**
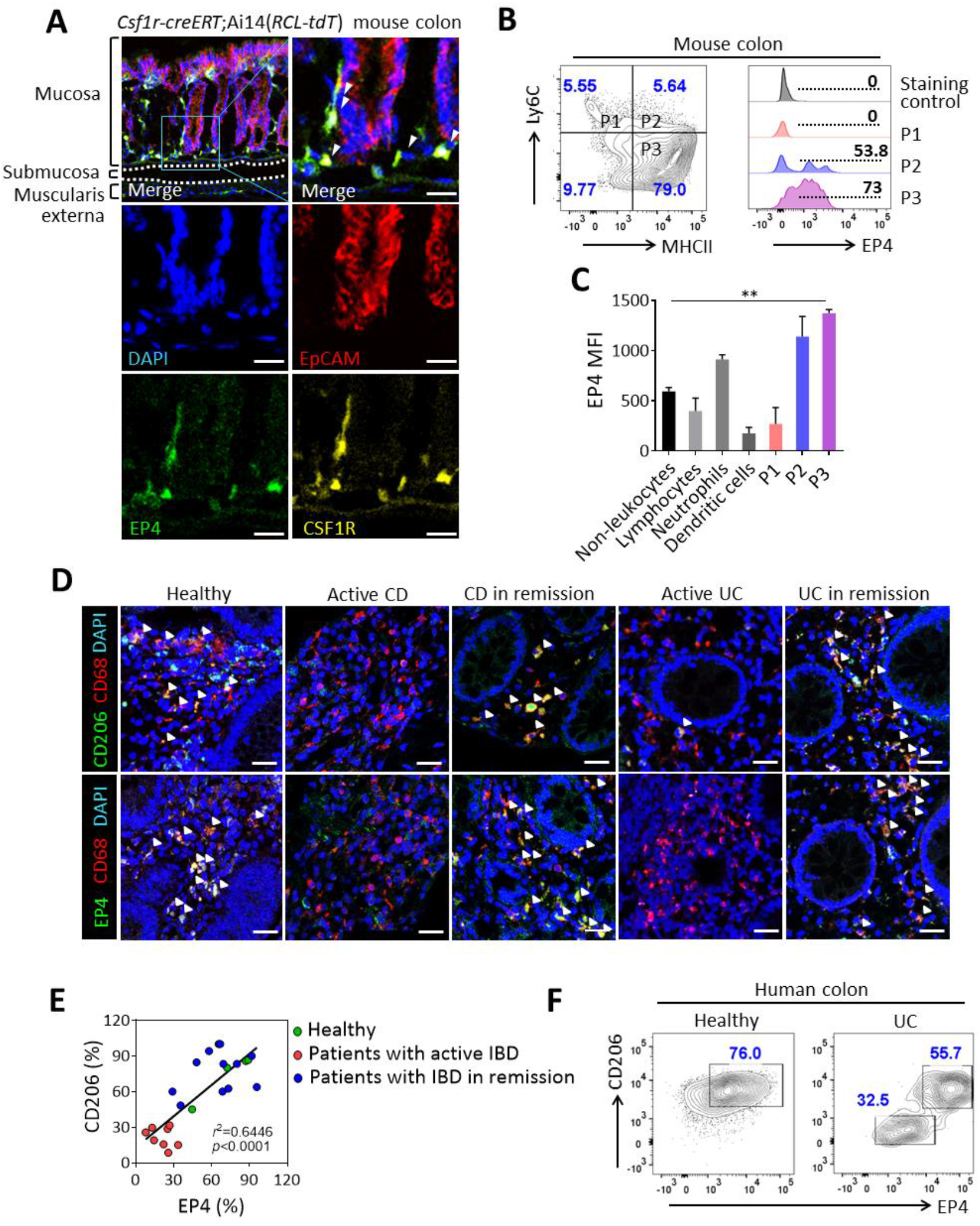
EP4 is expressed in mature intestinal macrophages and colocalized with CD206. (**A**) Confocal microscopy of colon tissue of tamoxifen-injected *Csf1r-creERT*;Ai14(*RCL-tdT*) mice stained for EP4 (green), CSF1R (yellow), EpCAM (red), and nuclei (DAPI; blue). Scale bar, 25 μm. Arrowheads indicate EP4^+^CSF1R^+^ cells. (**B**) Representative contour plot and histograms of the expression of EP4 in P1 (monocytes), P2 (intermediate macrophages), and P3 (mature macrophages) gated on a CD45^+^CD11b^+^Ly6G^−^CD64^+^ population. (**C**) Expression intensity of EP4 in non-leukocytes (CD45^−^), lymphocytes (CD45^+^CD11b^−^), neutrophils (CD45^+^CD11b^+^Ly6G^+^), dendritic cells (CD45^+^CD11b^+^Ly6G^−^Ly6C^−^CD64^−^), P1, P2, and P3. MFI, mean fluorescence intensity determined by flow cytometry. n = 4 mice per group. (**D**) Representative confocal microscopy of sectioned colon tissues of normal, active CD, CD in remission, active UC, and UC in remission stained for CD206 or EP4 (green), CD68 (red), and nuclei (DAPI; blue). Arrowheads indicate colocalization of EP4 or CD206 with CD68. Scale bar, 25 μm. (**E**) Percentages of populations each colocalized cell type in the lamina propria. *r*^2^ = 0.6446 analyzed by linear regression. *p*<0.0001. (**F**) Representative contour plots of EP4^+^CD206^+^ macrophages (gated on the CD45^+^CD11b^+^CD68^+^ population) in the mucosal layers of normal (n = 4) and patients with UC (n = 1) and CD (n=2). In panel C, statistical significance was determined by one-way ANOVA with Dunnett’s multiple comparisons test. \*\**P* < 0.01.

In line with the murine data, high expression of EP4 was found on CD68^+^ macrophages in the colonic mucosa of healthy individuals. Confocal microscopy of EP4 and CD206 revealed a positive correlation (*r*^2^ = 0.6446) of these markers on CD68^+^ macrophages in the healthy human colon (Fig. 1, D and E, Table S1). The percentage of EP4^+^ CD206^+^ macrophages was reduced in patients with active IBD (CD = 6, UC = 4) compared with healthy individuals. Interestingly, the percentage of EP4^+^ CD206^+^ macrophages was recovered in patients with IBD in remission (CD = 6, UC = 4) (Fig. 1E). In line, flow cytometric analysis of colonic biopsies from healthy individuals (n = 4) confirmed that EP4 is mainly expressed by CD206^+^ mucosal macrophages (76 %; Fig. 1F). In contrast, EP4^+^CD206^+^ macrophages were decreased to 55 % in patients suffering from active UC, together with the appearance of a double negative monocytic population (patient information is provided in Table S1).

### Deficiency of EP4 in macrophages impairs mucosal repair during intestinal inflammation

To evaluate the role of myeloid derived EP4 during intestinal inflammation, tamoxifen-inducible myeloid cell-specific EP4 knockout mice (*Csf1r*-*Ptger4*^−/−^), which were confirmed to have a specific gene recombination of *Ptger4* in CSF1R^+^ myeloid cells (Fig. S2, S3A-G), were used. Wildtype (WT) and *Csf1r*-*Ptger4*^−/−^ mice were exposed to 2.5% dextran sodium sulfate (DSS) in drinking water for 5 days (Fig. 2A). As compared to WT mice, which recovered 19 days post treatment (dpt) with DSS, *Csf1r*-*Ptger4*^−/−^ mice remained at 85% of their starting weight 19 dpt (Fig. 2A), which was correlated with a higher disease activity index, delayed recovery of symptoms and shorter colon length over time (Fig. 2, B and C). However, myeloid cell specific EP4 knockout was shown to induce a similar level of inflammation with no differences between the two groups based on myeloperoxidase (MPO) activity (Fig. S4A), infiltration of total leukocytes (Fig. S4B), neutrophils (Fig. S4C), and monocytes (Fig. S4D). Histological scores were similar during the early inflammatory phase, whereas *Csf1r*-*Ptger4*^−/−^ mice showed significantly higher scores during the late recovery phase due to crypt loss and erosion, indicating a defect in the epithelial repair process in these conditional knockout mice (Fig. 2D).

**Fig. 2.**
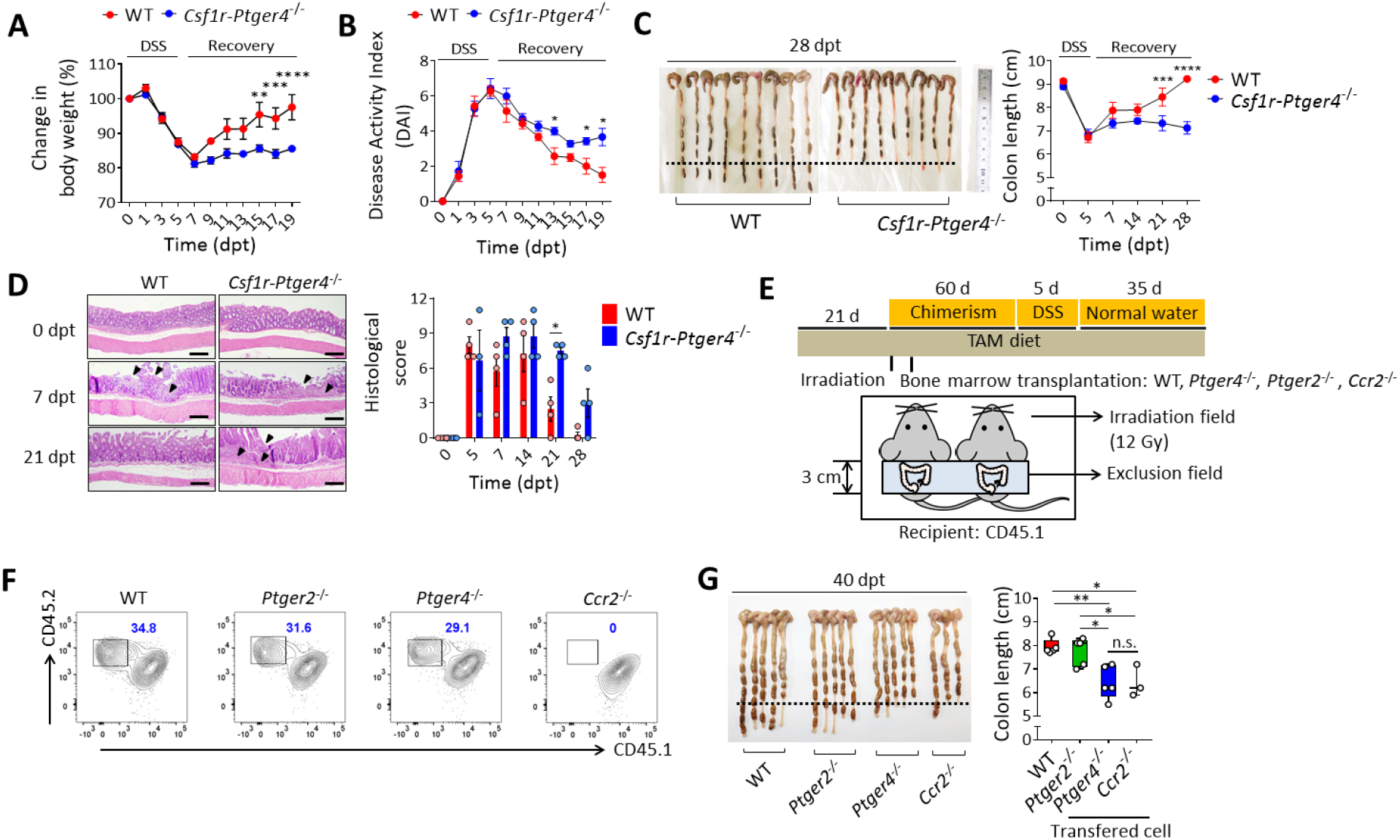
Monocyte derived EP4^+^ macrophages contribute to complete resolution of colitis. **A-D**, Wild-type and *Csf1r*-*Ptger4*^−/−^ mice were given water containing 2.5 % DSS (w/v) for 5 d, followed by regular water, and analyzed on indicated time points. dpt, days post treatment with DSS. (**A**) Weights are shown as a percentage of the initial weight of WT and KO mice. n = 7 mice. (**B**) Disease activity index, a composite measure of weight loss, stool consistency, and blood in stool, is shown. (**C**) Comparison of colon lengths. n = 9 mice at each time point. (**D**) Representative hematoxylin and eosin-stained sections of the colon. Arrowhead indicates erosion and crypt loss. Histological scores at each time point are shown at the right. **E-G**, Mixed bone marrow chimeras were generated by reconstituting CD45.1^+/+^ lethally irradiated recipients with WT, *Ptger4*^−/−^, *Ptger2*^−/−^, and *Ccr2*^−/−^ bone marrow. TAM diet; diet containing 400 mg/kg of tamoxifen. (**E**) Overview of experimental set-up for the generation of mixed bone marrow chimera. Irradiation was performed with shielding of the abdomen area. (**F**) Representative contour plots of reconstituted intestinal macrophages of mixed bone marrow chimeras. Data are representative of 1 of 2 independent experiments. (**G**) Comparison of colon lengths. In panels A-D, statistical significance was determined by two-way ANOVA. In panels G, one-way ANOVA was performed. \**P* < 0.05; \*\**P* < 0.01; \*\*\**P* < 0.001.

To examine whether EP4 in CSF1R^+^ macrophages contributes to intestinal repair, we adoptively transferred WT bone marrow derived macrophages, dendritic cells and neutrophils, the three major populations expressing CSF1R in the murine intestine (Fig. S2), into *Csf1r*-*Ptger4*^−/−^ recipient mice (Fig. S5A). As expected, transfer of WT macrophages into conditional KO mice, but not dendritic cells or neutrophils, significantly increased colon length after DSS colitis, indicating that EP4^+^ macrophages may play an important role in intestinal regeneration (Fig. S5B). To test whether these regeneration-inducing macrophages originated from blood monocytes, irradiated CD45.1 mice were transplanted with WT, prostaglandin E receptor 2 (*Ptger2;* EP2) KO, *Ptger4* (EP4) KO, or C-C motif chemokine receptor 2 (*Ccr2*) KO bone marrow cells, and subjected to DSS colitis (Fig. 2E-G). Of note, both mice receiving EP4 KO and CCR2 KO bone marrow cells had shorter colons when compared with WT mice (Fig.2G). In contrast, mice that received EP2 KO bone marrow cells did not show significant differences in colon length compared with WT controls (Fig. 2G). Therefore, both infiltration of blood monocytes and subsequent expression of EP4 might thus be necessary for promoting a wound healing phenotype in monocyte-derived intestinal macrophages. Collectively, these findings demonstrate that monocyte-derived EP4^+^ macrophages are required for mucosal repair during intestinal inflammation.

### EP4^+^ macrophages regulate intestinal crypt restoration and epithelial cell proliferation upon inflammatory damage

To investigate the epithelial defect in *Csf1r*-*Ptger4*^−/−^ mice during intestinal inflammation, we quantified the crypt numbers per unit length at 7 and 21 dpt. While the number of crypts decreased by 7 dpt in both WT and *Csf1r*-*Ptger4*^−/−^ mice, *Csf1r*-*Ptger4*^−/−^ mice had a significantly lower number of crypts compared with WT at 21 dpt (Fig. 3A). Furthermore, flow cytometric analysis of colonic epithelial cells (Fig. 3B) and immunohistochemistry on whole colons (Fig. 3C) showed that the number of Ki67^+^ proliferating epithelial cells was significantly lower in *Csf1r*-*Ptger4*^−/−^ mice than in WT mice by 21 dpt, pointing towards epithelial defects in the absence of myeloid derived EP4. To explore the epithelial response during the course of colitis, we identified the gene expression level of markers for crypts, including leucine rich repeat containing G protein-coupled receptor 5 (*Lgr5;* crypt-base columnar cells, CBCs (*22*)), Bmi1 proto-oncogene (*Bmi1*; reserve stem cells, RSCs (*23*)), and regenerating family member 4 ((*Reg4*)/lysozyme C-1 (*Lyz1*)/ lysozyme C-2 (*Lyz2*); deep crypt epithelial cells (*24*)) (Fig. 3D). All markers were decreased at 7 dpt, representing the loss of crypts in the inflammatory phase. From 14 dpt onwards, these genes were observed to be transcriptionally increased in WT mice, whereas their expression remained low in *Csf1r*-*Ptger4*^−/−^ mice. These results demonstrate that EP4^+^ macrophages might be required for the restoration of damaged crypts and for promoting epithelial cell proliferation.

**Fig. 3.**
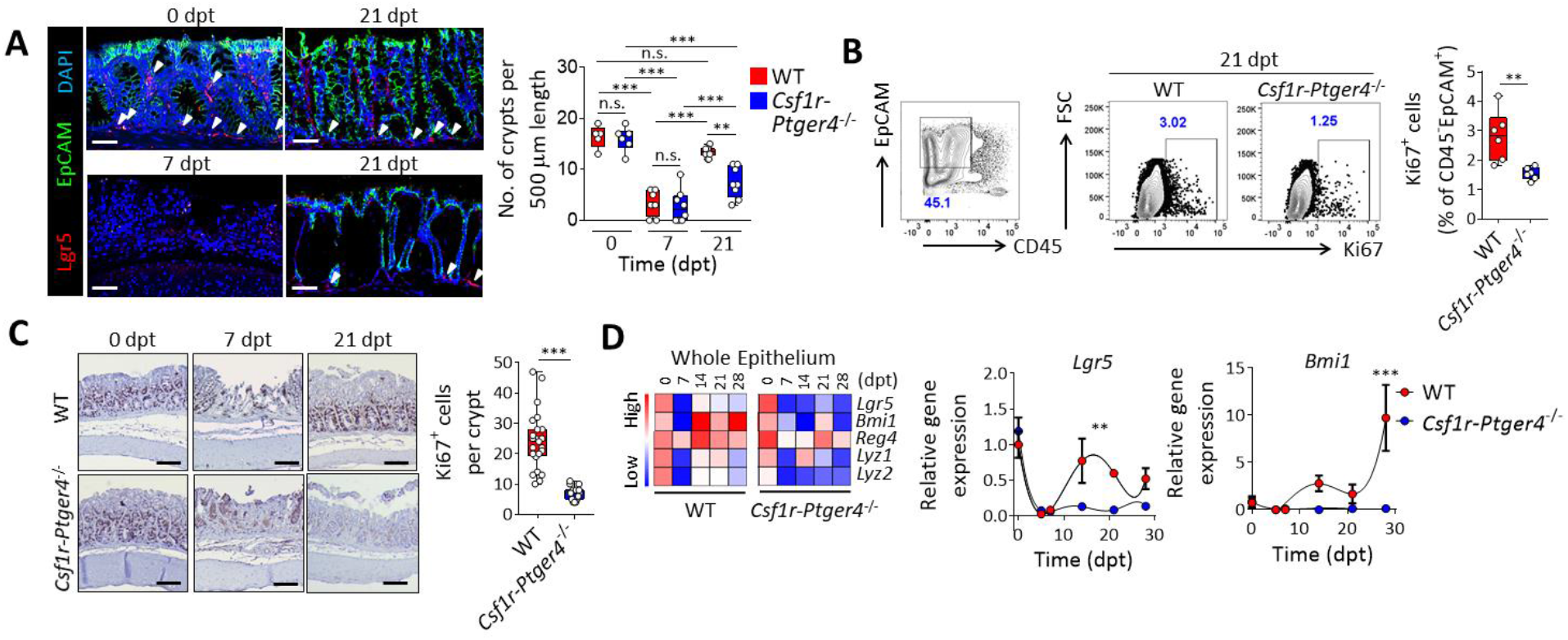
EP4 deficiency impairs the restoration of crypts and epithelial cell proliferation. **A-D**, WT and *Csf1r*-*Ptger4*^−/−^ mice were given 2.5 % DSS (w/v) for 5 d, followed by regular water, and analyzed at indicated time points. dpt, days post treatment with DSS. (**A**) Confocal images of colons from WT mice at 0, 7, and 21 dpt and from KO mice at 21 dpt stained for EpCAM (green), LGR5 (red), and DAPI (nuclei; blue) is shown at left. Scale bar, 25 μm. Quantitative number of crypts per 500 μm in colonic tissues are indicated on the right. Arrowheads indicate crypts having *Lgr5*^+^ cells. (**B**) Representative contour plots of Ki67^+^ epithelial cells (CD45^−^EpCAM^+^) at 21 dpt. Quantitative graph is shown on the right. (**C**) Representative image of Ki67 in colonic tissues from WT and KO mice at 0, 7, and 21 dpt. Number of Ki67^+^ cells per crypt are quantitated on the right. (**D**) Heatmap of time course of the gene expression for *Lgr5*, *Bmi1*, *Reg4*, *Lyz1*, and *Lyz2* in whole colonic epithelial cells isolated from WT and KO mice at indicated time points. The relative gene expression of *Lgr5* and *Bmi1* is shown on the right. n = 6 mice per time point. In panels A and D, statistical significance was determined by two-way ANOVA. In panels B and C, a two-tailed unpaired Student’s *t*-test was performed. \*\**P* < 0.01; \*\*\**P* < 0.001.

### EP4 signaling promotes the wound healing phenotype of colonic macrophages

After confirming that IL-4 was not an important factor for inducing wound healing macrophages in DSS colitis (Fig. S6, A-D), we next examined whether the expression of EP4 contributed to the development of wound healing macrophages during intestinal inflammation. Although colon shortening was observed in *Csf1r*-*Ptger4*^−/−^ (Fig. 2C), we observed no differences in the absolute numbers of macrophages between WT and KO mice during the entire experimental period (Fig. 4A). In contrast, a decrease in the number of CD206^+^ (Fig. 4B), as well as ARG1^+^ and RETNLA^+^ macrophages (Fig. 4C), was detected in *Csf1r*-*Ptger4*^−/−^ mice at 40 dpt. In addition, as shown in human colonic biopsies (Fig. 1F), we also detected a strong positive correlation between the expression of EP4 and CD206 in intestinal macrophages using flow cytometry (Fig. 4, D and E). Similarly, *in vitro* generated bone marrow derived macrophages (BMDMs) also required EP4 for their expression of CD206, ARG1, and RETNLA (Fig. S7, A and B). These results indicate the involvement of EP4 in inducing the expression of wound healing markers on macrophages, but not in the infiltration during intestinal inflammation.

**Fig. 4.**
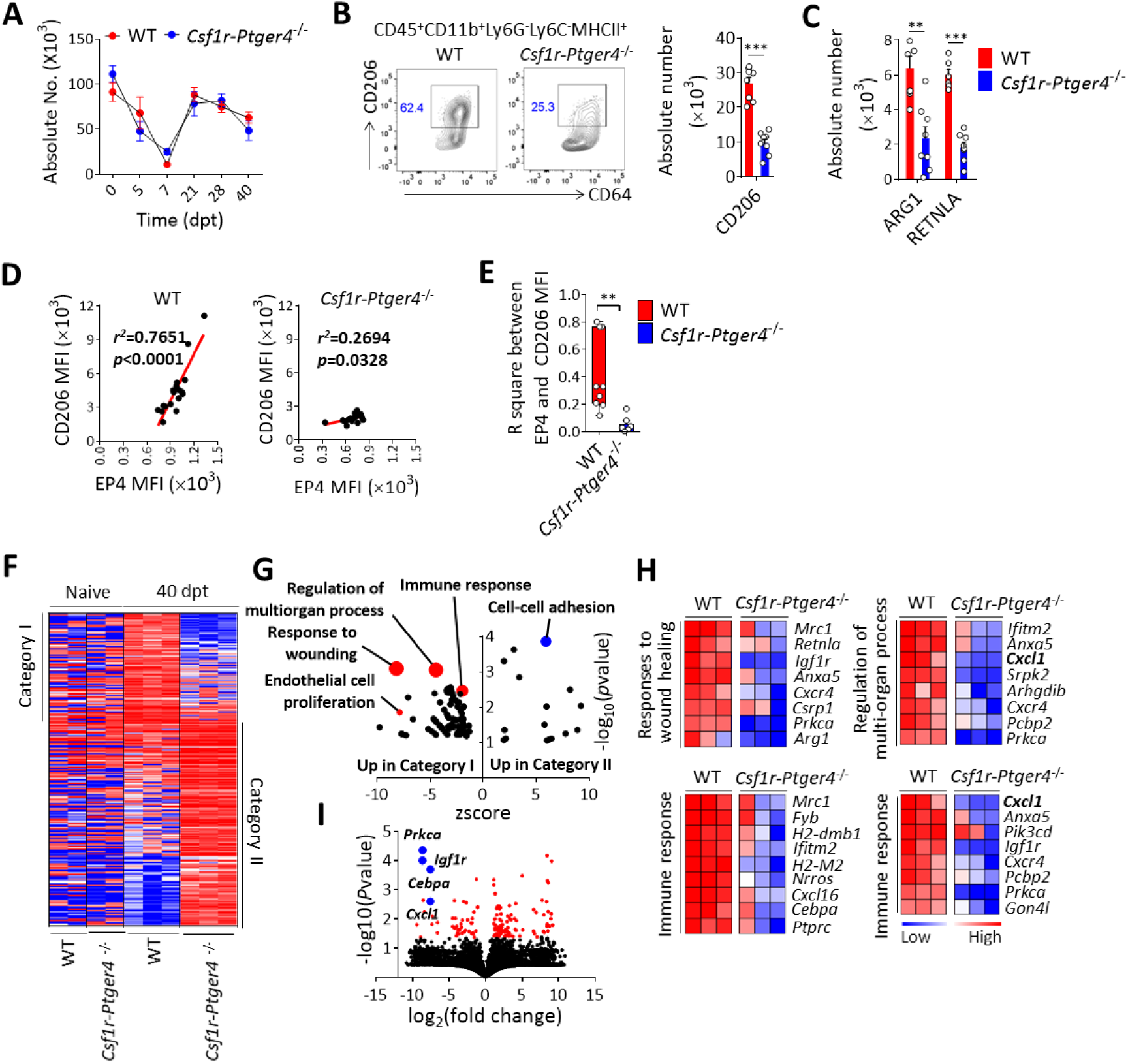
Wound healing phenotype of macrophages depends on the expression of EP4. **A-H**, WT and *Csf1r*-*Ptger4*^−/−^ mice were given 2.5 % DSS (w/v) for 5 d, followed by regular water, and analyzed at indicated time points. (**A**) Absolute number of mature colonic macrophages in WT and KO mice at indicated time points. (**B**) Representative contour plots of the CD206^+^ macrophages (gated on the CD45^+^CD11b^+^Ly6G^−^Ly6C^−^MHCII^+^ population) in the lamina propria of WT and KO mice at 40 dpt. Absolute number of CD206^+^ macrophages (x10^6^) of lamina propria cells is shown on the right. (**C**) Absolute number of ARG1^+^ and RETNLA^+^ macrophages in WT and KO mice at 40 dpt. (**D**) Correlation between the mean fluorescence intensity of EP4 and CD206 in WT and KO mice analyzed by flow cytometry. R squares in each mouse are indicated in (**E)**. MFI, mean fluorescence intensity. **F-I**, RNA-seq of colonic macrophages isolated from WT and KO mice at 0 and 40 dpt. (**F**) Heatmap depicts the significantly different genes between WT and KO at 40 dpt. (**G**) GO plot shows enriched gene ontologies in Category I (red circle) and Category II (blue circle). (**H**) Heatmaps depict the most differentially expressed genes in each GO. (**I**) Volcano plot showing differentially expressed genes between macrophages from WT and KO mice (red circle) at 40 dpt. Blue circles represent the most significantly downregulated genes in KO mice. In panels B, C, and E, a two-tailed unpaired Student’s *t*-test was performed. In panel D, linear regression was performed. \*\**P* < 0.01; \*\*\**P* < 0.001.

To define the mechanisms that underlie regeneration by EP4^+^ macrophages, we transcriptionally compared sorted WT and EP4^−/−^ macrophages isolated from the colon at 0 and 40 dpt by RNAseq. In total, 252 genes were differentially expressed (Fig. 4F) between WT and KO macrophages at 40 dpt (Fig. 4G). Using gene ontology, we observed a wound healing signature in WT macrophages, including mannose receptor C-type 1 (*Mrc1*), *Retnla*, insulin like growth factor 1 receptor (*Igf1r*), annexin A5 (*Anxa5*), C-X-C motif chemokine receptor 4 (*Cxcr4*), cysteine and glycine rich protein 1 (*Csrp1*), protein kinase C alpha (*Prkca*) and *Arg1*, which was reduced in KO macrophages. Concomitantly, the regulation of multiorgan processes and proliferation of endothelial cells were enriched in WT mice. Several genes involved in the immune response were also identified to be decreased in KO macrophages, including CCAAT enhancer binding protein alpha (*Cebpa*), *Cxcl1*, and *Prkca* (Fig. 4, H and I). Collectively, these results demonstrated that the expression of EP4 might oversee the overall features of wound healing macrophages.

### CXCL1 secretion by EP4^+^ macrophages is necessary for intestinal epithelial cell proliferation

To investigate whether soluble factors derived from EP4^+^ macrophages might stimulate epithelial repair, *Csf1r*-*Ptger4*^−/−^ mice were injected with supernatant of lamina propria mononuclear cells from the colon of WT mice (Fig. 5A). Injection of the supernatant resulted in increased colon length in *Csf1r*-*Ptger4*^−/−^ mice at 40 dpt, indicating that soluble factors were involved in epithelial repair (Fig. 5B). WNTs, previously identified molecules involving crypt regeneration as secreting factors from intestinal macrophages (*25*), were transcriptionally detected in low levels and showed no expression difference between WT and *Csf1r*-*Ptger4*^−/−^ intestinal macrophages (Fig. S8, A and B). Instead we identified CXCL1 as a potential factor in promoting epithelial regeneration (*26*). Interestingly, blocking CXCL1 with neutralizing antibodies in the supernatant before injection prevented epithelial repair during DSS colitis. In line, higher level of CXCL1 was detected in the supernatant of WT lamina propria mononuclear cells compared to *Csf1r*-*Ptger4*^−/−^ supernatant (Fig. 5C). CXCL1 RNA and protein levels were also decreased in whole colon lysates from *Csf1r*-*Ptger4*^−/−^ mice compared with WT mice at 40 dpt (Fig. 5D-E). Moreover, CXCL1 was predominantly localized in F4/80^+^ macrophages in the colonic lamina propria of WT mice (Fig. 5F) at 40 dpt as determined by IF staining, whereas *Csf1r*-*Ptger4*^−/−^ mice barely showed any expression of CXCL1 in the colon. These data indicated that macrophages might be the major producers of CXCL1 in the resolution phase of colitis and that the production of CXCL1 is dependent on EP4 signaling.

**Fig. 5.**
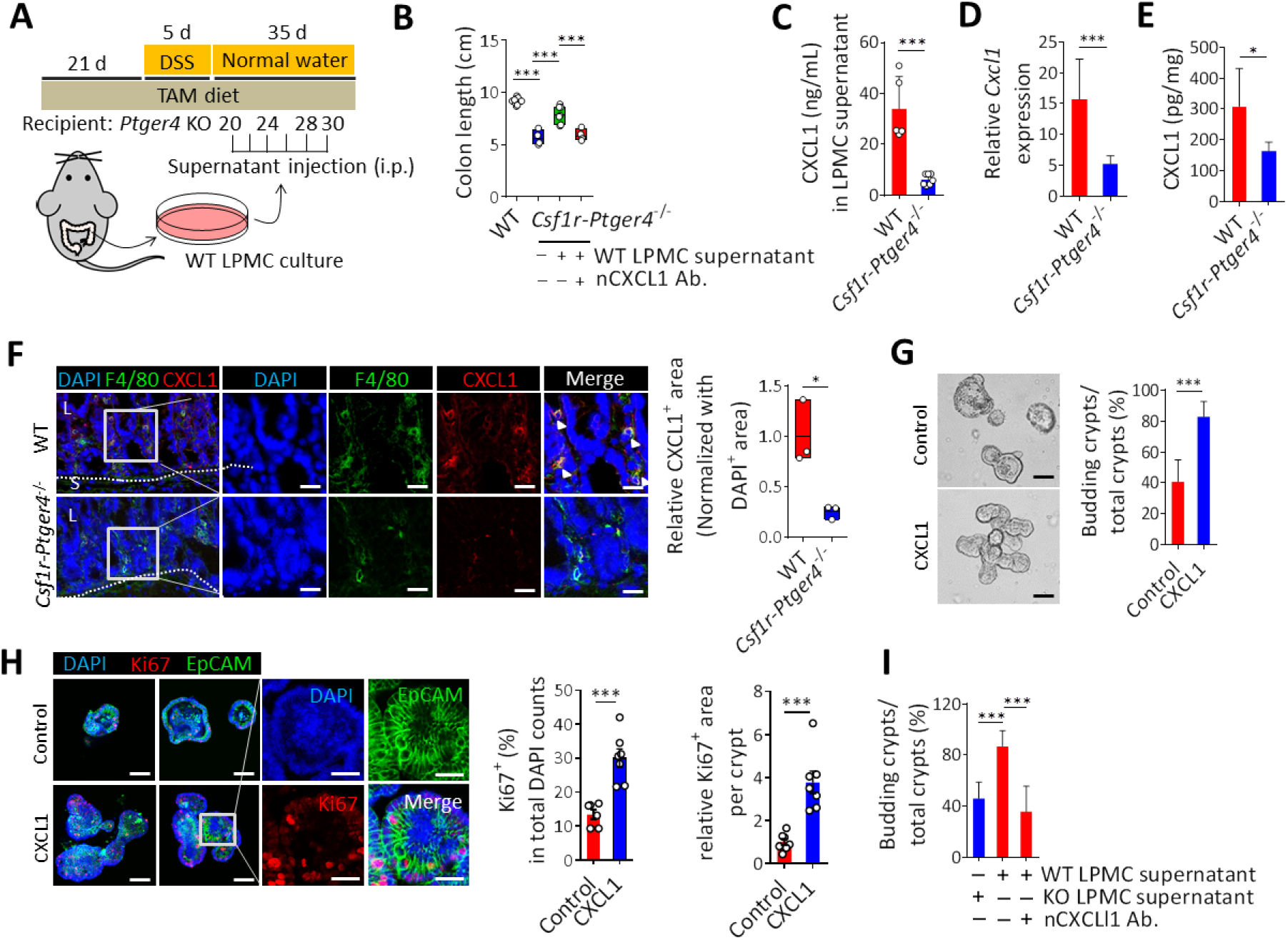
CXCL1 derived from EP4^+^ macrophages induces epithelial cell proliferation. **A-E**, WT and *Csf1r*-*Ptger4*^−/−^ mice were given 2.5 % DSS (w/v) for 5 d, followed by regular water, and analyzed at 40 dpt. Accordingly, 200 μL of culture supernatant of lamina propria mononuclear cells (LPMC) obtained from WT mice was intraperitoneally injected 6 times during the period of 20~30 dpt. Neutralizing antibodies for CXCL1 (20 μg/mL) was added to the supernatant before injection. (**A**) Overview of experimental set-up. (**B**) Comparison of colon lengths. (**C**) A total of 2 × 10^7^ LPMCs isolated from colons of WT and KO mice were cultured for 12 h. CXCL1 was determined in supernatant by enzyme-linked immunosorbent assay (ELISA). (**D**) mRNA expression of *Cxcl1* in whole colon lysates of WT and KO mice. (**E**) CXCL1 protein levels were determined in whole colon lysates of WT and KO mice. (**F**) Representative confocal images of WT and KO colons at 40 dpt stained for F4/80 (green), CXCL1 (red), and DAPI (nuclei; blue) at 40 dpt. Arrowheads indicate F4/80^+^CXCL1^+^ cells. Quantification of the CXCL1^+^ area was performed using the ImageJ and the CXCL1^+^ area was normalized relative to the DAPI^+^ area shown on the right. Scale bar, 25 μm. (**G**) WT organoids were cultured with or without 10 ng/mL of recombinant CXCL1 for 5 d (n = 3 independent cultures). Percentages of budding crypts relative to total crypts are shown on the right. Scale bar, 25 μm. (**H**) Ki67 and EpCAM staining in representative organoids cultured with 10 ng/mL of CXCL1. Percentages of Ki67^+^ cells per crypt and the relative Ki67^+^ area per crypt are shown on the right. Scale bar, 25 μm. (**I**) Organoids were treated with LPMC supernatant obtained from WT or KO mice for 5 d. Accordingly, 10 μg/mL of CXCL1 neutralizing antibody was added to the WT LPMC supernatant. Percentages of budding crypts relative to total crypts are shown on the right. In panels B and I, statistical significance was determined by one-way ANOVA. In panels C, D, E, F, G, and H, a two-tailed unpaired Student’s *t*-test was performed. \**P* < 0.05; \*\*\**P* < 0.001.

To evaluate the effect of macrophage-derived CXCL1 on epithelial cell function, we investigated it effect on murine intestinal organoids *in vitro*. Firstly, we confirmed that C-X-C motif chemokine receptor 2 (CXCR2), the receptor for CXCL1, was expressed on colonic epithelial cells (Fig. S9A), as well as by intestinal organoids (Fig. S9B) generated from mouse colons. To explore the direct effect of CXCL1 in the regeneration of epithelial crypts, we generated intestinal organoids with or without the addition of recombinant CXCL1. Notably, CXCL1 was shown to promote epithelial cell proliferation as determined by crypt budding efficiency (Fig. 5G), as well as the ratio of Ki67^+^ proliferating cells in the crypt (Fig. 5H). Next, we treated the regenerating crypts with supernatant of lamina propria mononuclear cells and confirmed higher budding efficiency in crypts co-cultured with supernatant of WT cells compared with those obtained from KO mice (Fig. 5I). Adding neutralizing antibodies for CXCL1 to the supernatant of WT cells reverted the budding efficiency to the level of KO samples. Collectively, these findings demonstrated that EP4^+^ macrophages stimulate epithelial cell proliferation in a CXCL1-dependent manner.

### Macrophages release CXCL1 via the PGE_2_/EP4/MAPKs signaling pathway

To define the exact molecular mechanisms of CXCL1 production by EP4^+^ macrophages, we first determined *Cxcl1* gene transcripts in WT BMDMs following stimulation with 10 μM of PGE_2_. At 48 h post treatment with PGE_2_, we observed that the *Cxcl1* mRNA was increased 30-fold (Fig. 6A). Notably, this was primarily mediated by EP4, as *Ptger4*^−/−^ BMDMs did not show increased production of CXCL1 upon treatment with PGE_2_, whereas *Ptger2*^−/−^ and WT BMDMs showed significantly increased production of CXCL1 (Fig. 6B, *P*<0.001). Treatment of WT BMDM supernatant preactivated with PGE_2_ led to the increased expression of *Ki67* in crypts. On the other hand, adding neutralizing antibodies for CXCL1 into WT supernatant abrogated this effect (Fig. 6C), supporting the stimulatory effect of budding by CXCL1 derived from PGE_2_-stimulated macrophages. In contrast, both CXCL1 and WT BMDM supernatant treatment resulted in decreased expression of *Lgr5, Reg4,* and *Lyz1* in organoids, suggesting a possible inhibition of the stemness of crypts while favoring epithelial differentiation and proliferation.

**Fig. 6.**
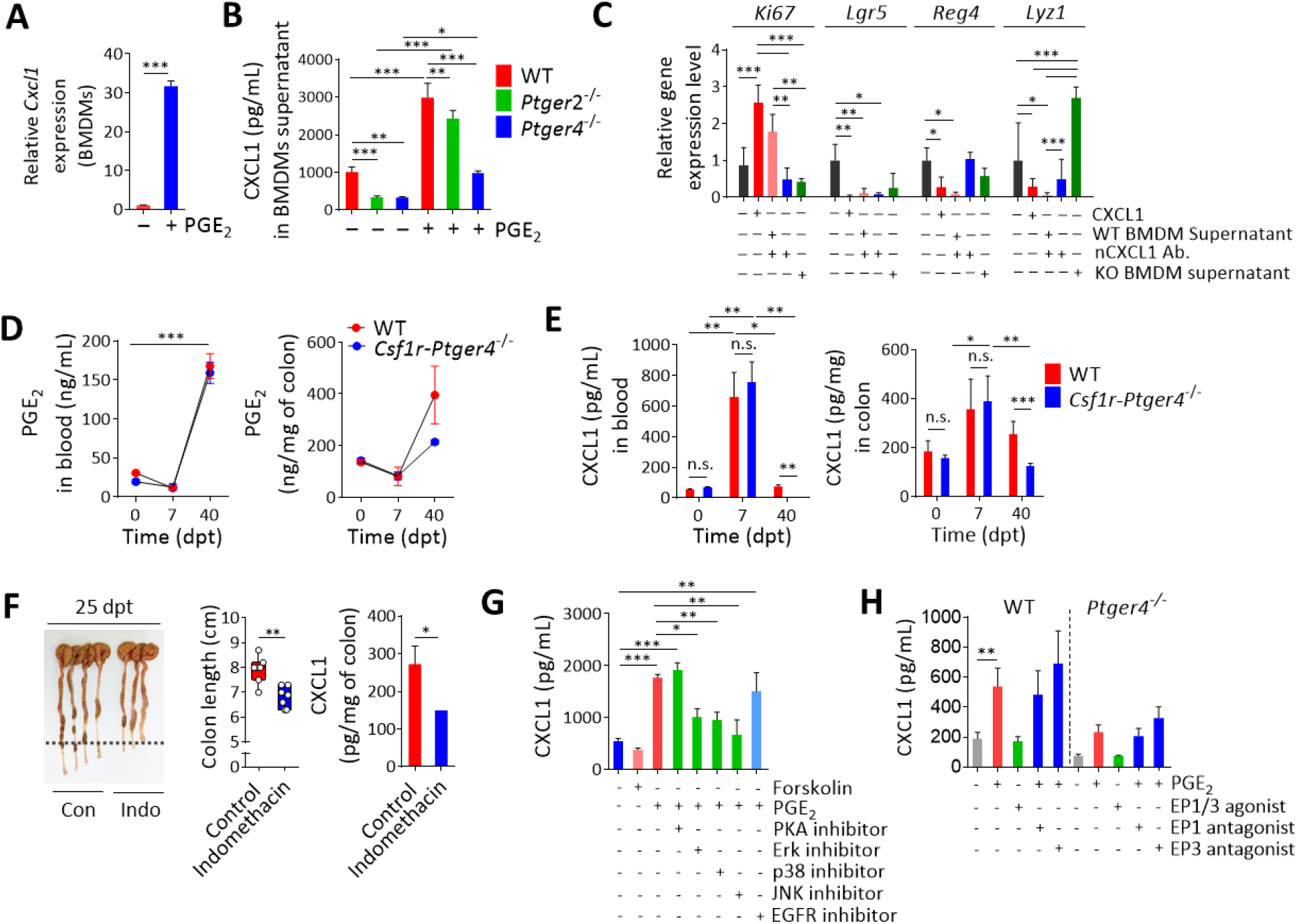
The PGE_2_/EP4/MAPKs signaling pathway control secretion of Cxcl1 in macrophages. (**A**) mRNA expression of *Cxcl1* in BMDMs treated with 10 μM of PGE_2_. (**B**) A total of 5 × 10^5^ BMDMs obtained from WT, *Ptger2*^−/−^ and *Ptger4*^−/−^ mice were cultured for 3 d in the presence or absence of PGE_2_. CXCL1 was determined in supernatant by ELISA. (**C**) Gene transcripts for *Ki67*, *Lgr5*, *Reg4*, and *Lyz1* of WT organoids were determined 2 d post treatment with 10 ng/mL CXCL1, WT BMDM sup, KO BMDM sup, 10 μg/mL of CXCL1 neutralizing antibody in WT BMDM supernatant. (**D**) The level of PGE_2_ was measured in blood and whole colon lysates of WT and KO mice at different time points during DSS colitis using ELISA. (**E**) CXCL1 was determined in the blood and whole colon lysates of WT and KO mice at different time points during DSS colitis. (**F**) Comparison of colon lengths (left and middle) and CXCL1 in whole colon lysates (right) in WT mice treated with indomethacin (10 mg/L in drinking water) from 15 until 25 dpt. (**G**) BMDMs were treated with forskolin (10 μM) or PGE_2_ in the presence of inhibitors for PKA (H89, 20 μM), ERK (PD98059, 1 μM), p38 (SB203580, 0.1 μM), JNK (SP600125, 0.5 μM), or EGFR (erlotinib, 20 μM) and incubated for 3 d before measuring CXCL1 in the supernatant. (**H**) BMDMs were obtained from WT and KO mice and treated with PGE_2_ in the presence of EP1 (SC-50189, 10 μM) or EP3 (L-798106, 10 μM) antagonist. The EP1/3 agonist, sulprostone, was added at a concentration of 10 μM. After 3 d, the level of CXCL1 in the supernatant was measured. In panels B, C, G, and H, statistical significance was determined by one-way ANOVA. In panels D and E, two-way ANOVA was performed. In panels A and F, a two-tailed unpaired Student’s *t*-test was performed. \**P* < 0.05; **P < 0.01; \*\*\**P* < 0.001.

We next analyzed the levels of PGE_2_ during DSS colitis to examine the possible correlation between PGE_2_ and CXCL1 *in vivo*. Interestingly, PGE_2_ was slightly decreased in the acute inflammatory phase while increased above homeostatic level during the recovery phase both in the blood and colon lysates of WT and *Csf1r*-*Ptger4*^−/−^ mice (Fig. 6D). In line with our hypothesis, the level of CXCL1 was lower in the blood and colon lysates derived from *Csf1r*-*Ptger4*^−/−^ mice at 40 dpt compared with those from WT mice (Fig. 6E). Blocking the endogenous activity of COX by administrating indomethacin during the last 10 days of the recovery phase also led to reduced colon lengths and CXCL1 levels in colon lysates (Fig. 6F).

To elucidate the mechanisms controlling the production of CXCL1 by EP4^+^ macrophages, BMDMs were pre-incubated with chemical activator or inhibitors for different pathways known to be regulated by PGE_2_ (Fig. 6G). We found that forskolin, an adenylyl cyclase activator, did not increase the level of CXCL1, indicating that intracellular cAMP accumulation does not mediate the production of CXCL1 in macrophages. Inhibition of a major EP4 downstream signaling molecule, protein kinase A (PKA), with H89 or the epidermal growth factor receptor (EGFR) inhibitor erlotinib did not inhibit the production of CXCL1 by PGE_2_-stimulated macrophages, suggesting that another signaling pathway other than EP4/cAMP/PKA or EGF signaling is involved in the upregulation of CXCL1 in macrophages by PGE_2_.

On the other hand, inhibition of the mitogen-activated protein kinase (MAPK) signaling pathway with ERK (PD98059), p38 (SB203580), and JNK (SP600125) inhibitors was able to significantly decrease the production of CXCL1 in PGE_2_ activated macrophages. Finally, we were able to show that EP1 and EP3 signaling do not play any role in the PGE_2_-mediated CXCL1 production in BMDMs using specific agonist and antagonist such as sulprostone (EP1/3 agonist), SC-50189 (EP1 antagonist) and L-798106 (EP3 (antagonist) (Fig. 6H). These observations suggested that PGE_2_ can activate MAPKs to induce synthesis of CXCL1 in macrophages through the ligation of EP4, but PKA, PKC, and EGFR are not involved in the signaling cascade.

### Liposomes macrophage-specific activation of MAPKs induces intestinal epithelial repair in EP4-deficient mice

To test whether activation of the MAPK-Cxcl1 pathway in macrophages could stimulate intestinal repair even in *Csf1r*-*Ptger4*^−/−^ mice we decided to use liposomes loaded with asiatic acid (AA), a well-known p38 MAPK activator (*27*) (Fig. 7A). Treatment with AA increased the production of CXCL1 in BMDMs, which was abrogated with preincubation with a p38 inhibitor (Fig. 7B). Next, we examined the therapeutic potential of AA in WT mice and observed that mice receiving AA had significantly increased colon lengths at 35 dpt compared to the vehicle group (Fig. 7C). Considering that MAPKs act as complex signaling molecules in a wide variety of cell types, we made AA-liposomes to specifically deliver drugs into macrophages *in vivo* (Fig. 7D). Accordingly, *Csf1r*-*Ptger4*^−/−^ mice injected intravenously with FNR-conjugated, AA-containing liposomes during the resolution phase (20~35 dpt) showed that FNR^+^ liposomes were mostly detected in F4/80^+^ macrophages in the lamina propria of the colon (Fig. 7E), and were distributed mostly in CD206^+^ or CD206^−^ intestinal macrophages (Fig. 7F). Interestingly, *Csf1r*-*Ptger4*^−/−^ mice receiving AA-liposomes showed longer colons compared with *Csf1r*-*Ptger4*^−/−^ mice receiving control liposomes (Fig. 7G). Administration of AA-liposomes was associated with increased proliferation of epithelial cells (Ki67^+^) in the mucosa of *Csf1r*-*Ptger4*^−/−^ mice (Fig. 6H) and increased expression of CXCL1 in F4/80^+^ macrophages (Fig. 7I).

**Fig. 7.**
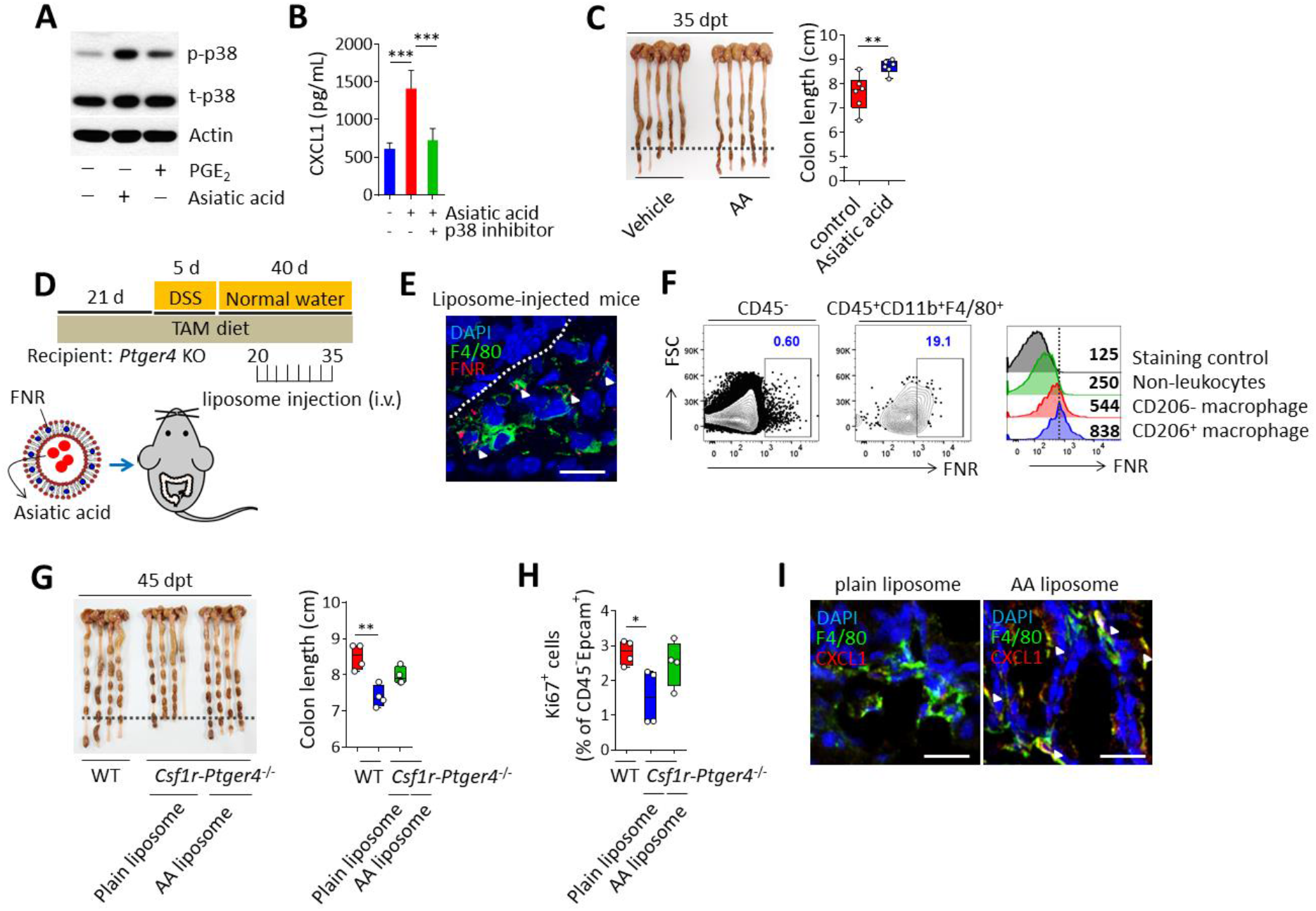
Therapeutic activation of MAPKs rescues intestinal regeneration of EP4-deficient mice. (**A**) Expression of phospho-p38 determined by western blotting. BMDMs were treated with 20 μM of asiatic acid (AA) or 10 μM of PGE_2_ for 1 h. (**B**) Determination of CXCL1 in supernatant was done by ELISA. BMDMs were treated with AA in the presence of a p38 inhibitor and cultivated for 3 d. (**C**) Comparison of colon lengths of WT mice (35 dpt) intraperitoneally injected with vehicle or AA (50 mg/kg) 8 times from 15 until 30 dpt during DSS-induced colitis. **D-I**, WT and *Csf1r*-*Ptger4*^−/−^ mice were given 2.5 % DSS (w/v) for 5 d, followed by regular water, and analyzed at 45 dpt. FNR-conjugated, asiatic acid-containing liposomes were iv injected for a total of 8 times from 20 until 35 dpt. (**D**) Overview of experimental set-up. (**E**) Confocal image of colon of a WT mouse at 2 d post liposome injection stained for F4/80 (green), FNR (red), and DAPI (nuclei; blue) (left). (**F**) Contour plots show FNR^+^ macrophages gated on CD45^+^CD11b^+^F4/80^+^ cells compared with CD45^−^ cells. Histograms show FNR intensities of CD45^−^ non-leukocytes, CD206^−^ macrophages, and CD206^+^ macrophages (right). Scale bar, 25 μm. (**G**) Comparison of colon lengths of WT and KO mice injected with plain liposomes or loaded with AA. (**H**) Percentages of Ki67^+^ epithelial cells in total CD45^−^EpCAM^+^ cells are shown. (**I**) Confocal images of colons of KO mice injected with plain liposomes or loaded with AA at 45 dpt stained for F4/80 (green), CXCL1 (red), and DAPI (nuclei; blue) (left). All data are representative of 2 independent experiments. In panels B, G, and H, statistical significance was determined by one-way ANOVA. In panel C, a two-tailed unpaired Student’s *t*-test was performed. \**P* < 0.05; \*\**P* < 0.01; \*\*\**P* < 0.001.

## Discussion

In this study, we identified EP4^+^ macrophages as the major drivers of epithelial regeneration during recovery from intestinal inflammation. Our data indicate that PGE_2_ via the EP4 promote the differentiation of a pro-resolving tissue protective monocyte-derived macrophages co expressing CD206. EP4^+^ CD206^+^ macrophages orchestrate the differentiation and proliferation of epithelial progenitors into their descendants during the resolution phase of inflammation. Under the influence of PGE_2_, activation of the MAPKs signaling pathway in EP4^+^ macrophages induced the secretion of CXCL1 inducing epithelial cell proliferation *in vivo* and in intestinal organoids. Taking advantage of the phagocytic capability of macrophages, we have demonstrated the possibility of generating a macrophage-targeted drug delivery system to enhance MAPKs signaling and potentiate macrophages mucosal healing phenotype. Overall, for the first time we report that macrophage-derived CXCL1 is essential for mucosal healing and may represent a novel therapeutic target to promote remission in patient suffering from IBD.

For its functional homeostasis the intestinal epithelial barrier relies on constant replenishment of enterocytes differentiating from stem cells residing at the bottom of the mucosal crypts (*28*). Self-renewal of the intestinal stem cells is therefore essential for mucosal homeostasis and it is therefore protected by possible injury such as radiation and inflammation (*10, 29–31*). In the case of acute radiation injury, release of Wnt signaling molecules by macrophages is essential to protect the colonic epithelial stem cell niche and consequent restoration of intestinal homoeostasis. It should be noted that WT and EP4 null intestinal macrophages have no differences on their expression of *Wnt* transcripts in our RNAseq suggesting no role for PGE_2_ on *Wnt* induction in macrophages (Fig. S6). In line, expression levels of *Wnt* were much lower (average normalized log2 = 0.428) when compared to those of *Cxcl1* during mucosal healing (average normalized log2 = 9.841) in intestinal macrophages. It might be possible that Wnts are mainly secreted during homeostasis and during acute inflammation, whereas CXCL1 seems preferentially expressed during the recovery phase by fully differentiated mature macrophages (CD45^+^CD11b^+^Ly6G^−^CD64^+^Ly6C^−^MHCII^+^ cells). Probably there is a division of labor between Wnt mainly involved in protection of stem cells during an inflammatory phase, while CXCL1 maybe be involved in the induction of epithelia proliferation and differentiation accelerating epithelial regeneration leading to mucosal healing. It would be of interest to define if CXCL1 also relay on Yap, a downstream transcriptional effector of Hippo signaling in promoting intestinal stem cells survival and inducing a regenerative program via the Egf pathway (*32*). In line, we found that recombinant CXCL1 stimulated the differentiation and proliferation of spheroids to budding organoids while reducing the expression of genes related to stemness, implicating CXCL1 in the reprogramming of crypts and in driving active mucosal healing. Ensuring adequate crypt budding by CXCL1 might potentiate intestinal epithelial restoration, enabling restoration of gene transcripts for stem cell markers in the whole epithelium. Our finding is supported by previous evidences showing a delayed skin wound healing in mice deficient for the CXCL1 receptor CXCR2 (*33*). Thus, supporting the idea that the PGE_2_/EP4/CXCL1 axis in macrophages is a novel yet critical player of mucosal healing. Although we found phosphorylation of CXCR2 on epithelial cells in organoids following CXCL1 treatment (Fig. S8B), it would be interesting to investigate whether the responses to CXCL1 may vary in different cell types of the intestinal stem cell niche.

Even during homeostasis, the gut has typically higher PGE_2_ levels compared to other tissue such as bone marrow and liver (*20*). However, there is conflicting evidence regarding the level of PGE_2_ during intestinal inflammation. Peng *et al*. showed that PGE_2_ was significantly decreased in the colon of patients with UC, as well as in DSS colitis (*34*), while Maseda *et al.* showed increased colonic PGE_2_ during T cell transfer colitis (*35*). Using a model of DSS colitis, we observed a slight decrease of PGE_2_ during the acute inflammatory phase with a clear increase of PGE_2_ both in the colon and blood during the late phase of resolution of inflammation. This is in line with a previous study showing increased PGE_2_ in the resolution phase of zymosan-induced peritonitis (*25*). Although the exact source of intestinal PGE_2_ during inflammation remains to be determined, these results suggest that the production of PGE_2_ is increased during tissue repair and maybe necessary for mucosal healing. One possible candidate for the production of PGE_2_ could be the mesenchymal stem cells (MSCs), as shown in a colonic mucosal excision model (*36*). In fact, MSCs are known to induce alternatively activated macrophages via the production of PGE_2_ (*37–39*), suggesting their potential involvement in the stimulation of macrophage differentiation to support the crypt regeneration.

Genome-wide association studies have previously correlated SNPs in the proximity of the *PTGER4* gene region with a worsen disease outcome in patients with CD (*21*). Here in line, we showed that EP4 mRNA expression was significantly downregulated in patient with active CD and UC, but expression was restored in UC patient in remission. Therefore, our study revealed an essential role of EP4 in the differentiation of wound healing macrophages in the intestine, which can be targeted as a new treatment strategy for IBD. Future work on the expression of EP4 and CD206 in macrophages based on the genetic information together with tracking long-term therapeutic responses might provide further valuable information to develop patient specific therapeutic options.

The presence of CD206^+^ regulatory macrophages has been proposed as a favorable prognostic factor in determining the response rate to anti-TNFα therapy in patients with CD (*15, 16*). In addition, recent studies even suggest that induction of CD206^+^ macrophages vai the Fc portion of ipilimumab mediating the activation of the Fcγ receptor maybe an essential component of the anti-TNFα therapy (*40*). The correlation between the expression of EP4 and CD206 observed in our study indicates a new mechanistic role of PGE_2_ in inducing wound healing macrophages.

Although additional work is needed to understand the mechanisms responsible for resolution of intestinal inflammation, accumulation of EP4^+^ CD206^+^ pro-resolving macrophages in the intestinal microenvironment seems to be crucial to re-establish gut tissue homeostasis. The identification of environmental cues such as PGE_2_ regulating intestinal macrophage phenotypes and functions have deepened our understanding of their essential role in homeostasis and in inflammation. Based on these findings, therapeutic targeting of macrophages represents a novel appealing strategy to re-educate the intestinal immune microenvironment and restore tissue homeostasis after inflammation.

## Acknowledgments

We would like to thank Shuh Narumiya (Kyoto University Graduate School of Medicine, Japan) for providing B6.129S6(D2)-*Ptger4*^*tm1.1Matb*^/BreyJ (EP4 floxed) mice, and Jae Hoon Choi (Hanyang University, South Korea) for B6.129S4-*Ccr2^tm1Ifc^*/J mice (*Ccr2*^−/−^).

## Author contributions

Y.N. and S.S. conceptualized this study; Y.N. and D.J. designed the experiments; Y.N., D.J., G.G., M.J., H.R., S.S., and J.P. performed the experiments; Y.N., D.J., H.H., B.P., S.L., H.L. and I.C. analyzed data; Y.N. and D.J. interpreted data; H.K., Y.K., T.S., S.R., K.P., J.I., and Y.L. provided vital reagents; Y.N., D.J., M.S., G.M., and S.S. contributed to manuscript writing; Y.N. wrote the manuscript.

## Data and materials availability

Extended data will be available after publication.

**Correspondence and requests for materials** should be addressed to S.S. or G.M. Peer review information (please complete this sentence). Reprints and permissions information is available at (please complete this sentence).

## Supplementary Materials

Materials and Methods and Figures S1-S8.

