## Supplemental info for "E prostanoid receptor 4 expressing macrophages promote the regeneration of the intestinal epithelial barrier upon inflammation"

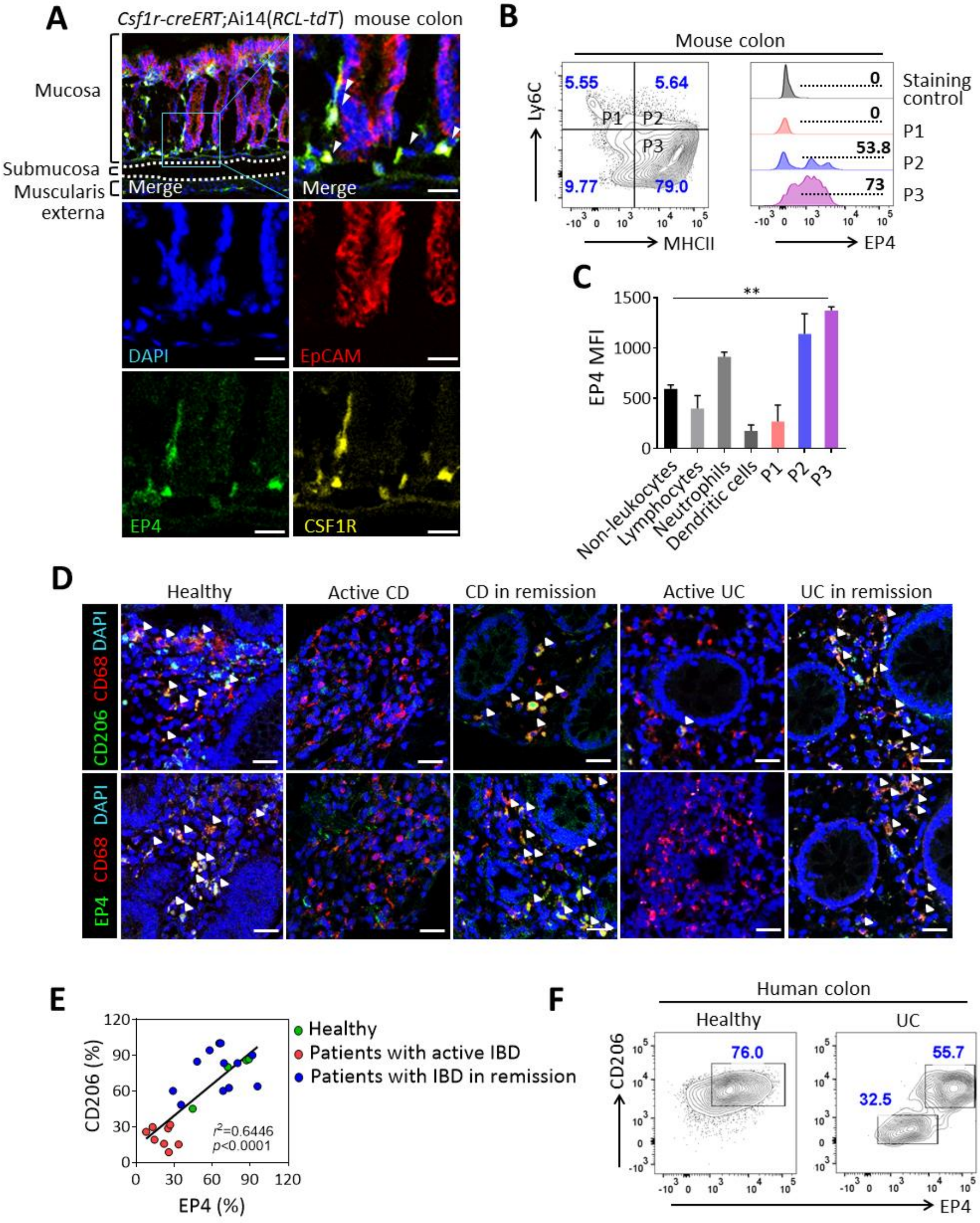


**Fig. 1. EP4 is expressed in mature intestinal macrophages and colocalized with CD206.** (**A**) Confocal microscopy of colon tissue of tamoxifen-injected *Csf1r-creERT*;Ai14(*RCL-tdT*) mice stained for EP4 (green), CSF1R (yellow), EpCAM (red), and nuclei (DAPI; blue). Scale bar, 25 µm. Arrowheads indicate EP4$^+$CSF1R$^+$ cells. (**B**) Representative contour plot and histograms of the

expression of EP4 in P1 (monocytes), P2 (intermediate macrophages), and P3 (mature macrophages) gated on a $CD45^+CD11b^+Ly6G^-CD64^+$ population. (**C**) Expression intensity of EP4 in non-leukocytes ($CD45^-$), lymphocytes ($CD45^+CD11b^-$), neutrophils ($CD45^+CD11b^+Ly6G^+$), dendritic cells ($CD45^+CD11b^+Ly6G^-Ly6C^-CD64^-$), P1, P2, and P3. MFI, mean fluorescence intensity determined by flow cytometry. n = 4 mice per group. (**D**) Representative confocal microscopy of sectioned colon tissues of normal, active CD, CD in remission, active UC, and UC in remission stained for CD206 or EP4 (green), CD68 (red), and nuclei (DAPI; blue). Arrowheads indicate colocalization of EP4 or CD206 with CD68. Scale bar, 25 µm. (**E**) Percentages of populations each colocalized cell type in the lamina propria. $r^2 = 0.6446$ analyzed by linear regression. $p<0.0001$. (**F**) Representative contour plots of $EP4^+CD206^+$ macrophages (gated on the $CD45^+CD11b^+CD68^+$ population) in the mucosal layers of normal (n = 4) and patients with UC (n = 1) and CD (n=2). In panel C, statistical significance was determined by one-way ANOVA with Dunnett's multiple comparisons test. $^{**}P < 0.01$.

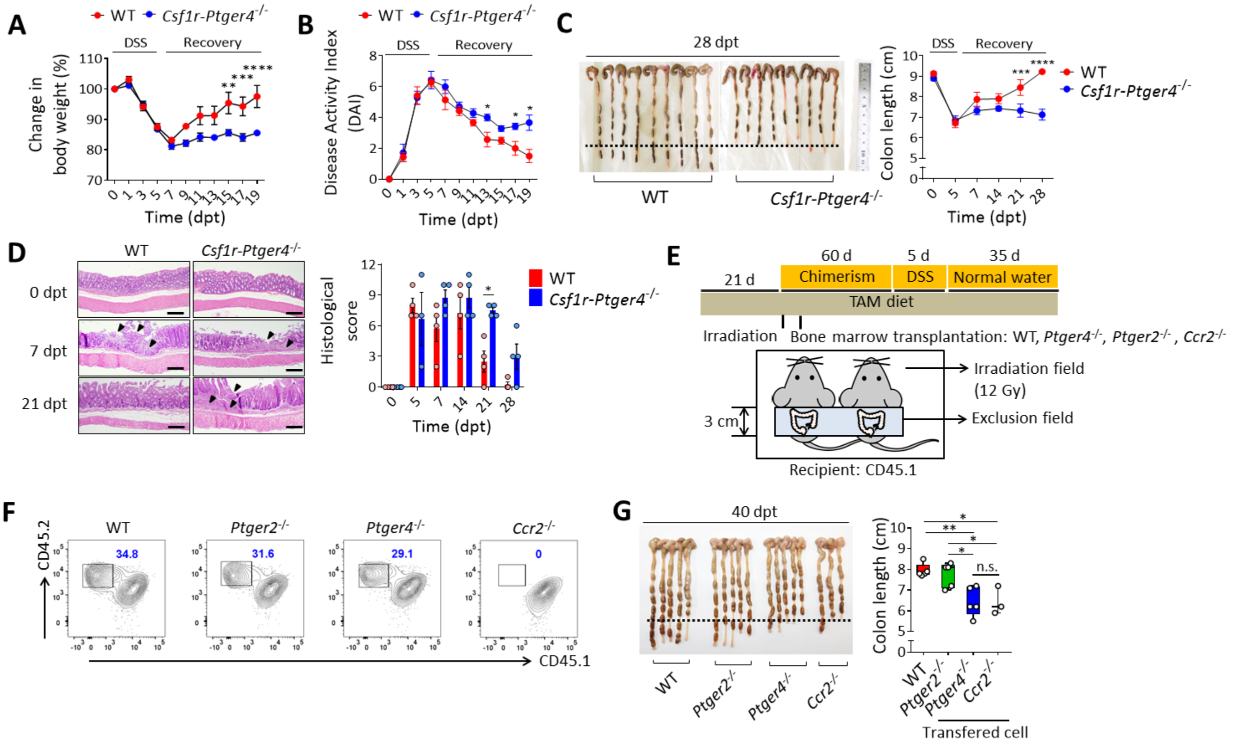


**Fig. 2. Monocyte derived EP4$^{+}$ macrophages contribute to complete resolution of colitis. A-D**, Wild-type and *Csf1r-Ptger4*$^{-/-}$ mice were given water containing 2.5 % DSS (w/v) for 5 d, followed by regular water, and analyzed on indicated time points. dpt, days post treatment with DSS. (**A**) Weights are shown as a percentage of the initial weight of WT and KO mice. n = 7 mice. (**B**) Disease activity index, a composite measure of weight loss, stool consistency, and blood in stool, is shown. (**C**) Comparison of colon lengths. n = 9 mice at each time point. (**D**) Representative hematoxylin and eosin-stained sections of the colon. Arrowhead indicates erosion and crypt loss. Histological scores at each time point are shown at the right. **E-G**, Mixed bone marrow chimeras were generated by reconstituting CD45.1$^{+/+}$ lethally irradiated recipients with WT, *Ptger4*$^{-/-}$, *Ptger2*$^{-/-}$, and *Ccr2*$^{-/-}$ bone marrow. TAM diet; diet containing 400 mg/kg of tamoxifen. (**E**) Overview of experimental set-up for the generation of mixed bone marrow chimera. Irradiation

was performed with shielding of the abdomen area. (**F**) Representative contour plots of reconstituted intestinal macrophages of mixed bone marrow chimeras.

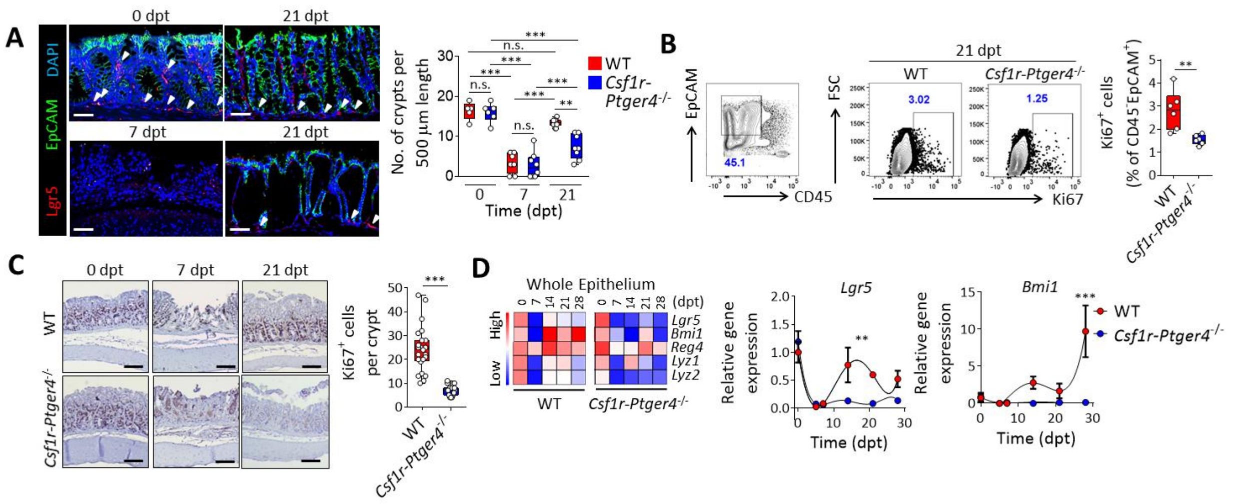


**Fig. 3. EP4 deficiency impairs the restoration of crypts and epithelial cell proliferation. A-D**, WT and *Csf1r-Ptger4*$^{-/-}$ mice were given 2.5 % DSS (w/v) for 5 d, followed by regular water, and analyzed at indicated time points. dpt, days post treatment with DSS. (**A**) Confocal images of colons from WT mice at 0, 7, and 21 dpt and from KO mice at 21 dpt stained for EpCAM (green), LGR5 (red), and DAPI (nuclei; blue) is shown at left. Scale bar, 25 µm. Quantitative number of crypts per 500 µm in colonic tissues are indicated on the right. Arrowheads indicate crypts having *Lgr5*$^{+}$ cells. (**B**) Representative contour plots of Ki67$^{+}$ epithelial cells (CD45$^{-}$EpCAM$^{+}$) at 21 dpt. Quantitative graph is shown on the right. (**C**) Representative image of Ki67 in colonic tissues from WT and KO mice at 0, 7, and 21 dpt. Number of Ki67$^{+}$ cells per crypt are quantitated on the right. (**D**) Heatmap of time course of the gene expression for *Lgr5*, *Bmi1*, *Reg4*, *Lyz1*, and *Lyz2* in whole colonic epithelial cells isolated from WT and KO mice at indicated time points. The relative gene expression of *Lgr5* and *Bmi1* is shown on the right. n = 6 mice per time point. In panels A and D, statistical significance was determined by two-way ANOVA. In panels B and C, a two-tailed unpaired Student's *t*-test was performed. $^{**}P < 0.01$; $^{***}P < 0.001$.

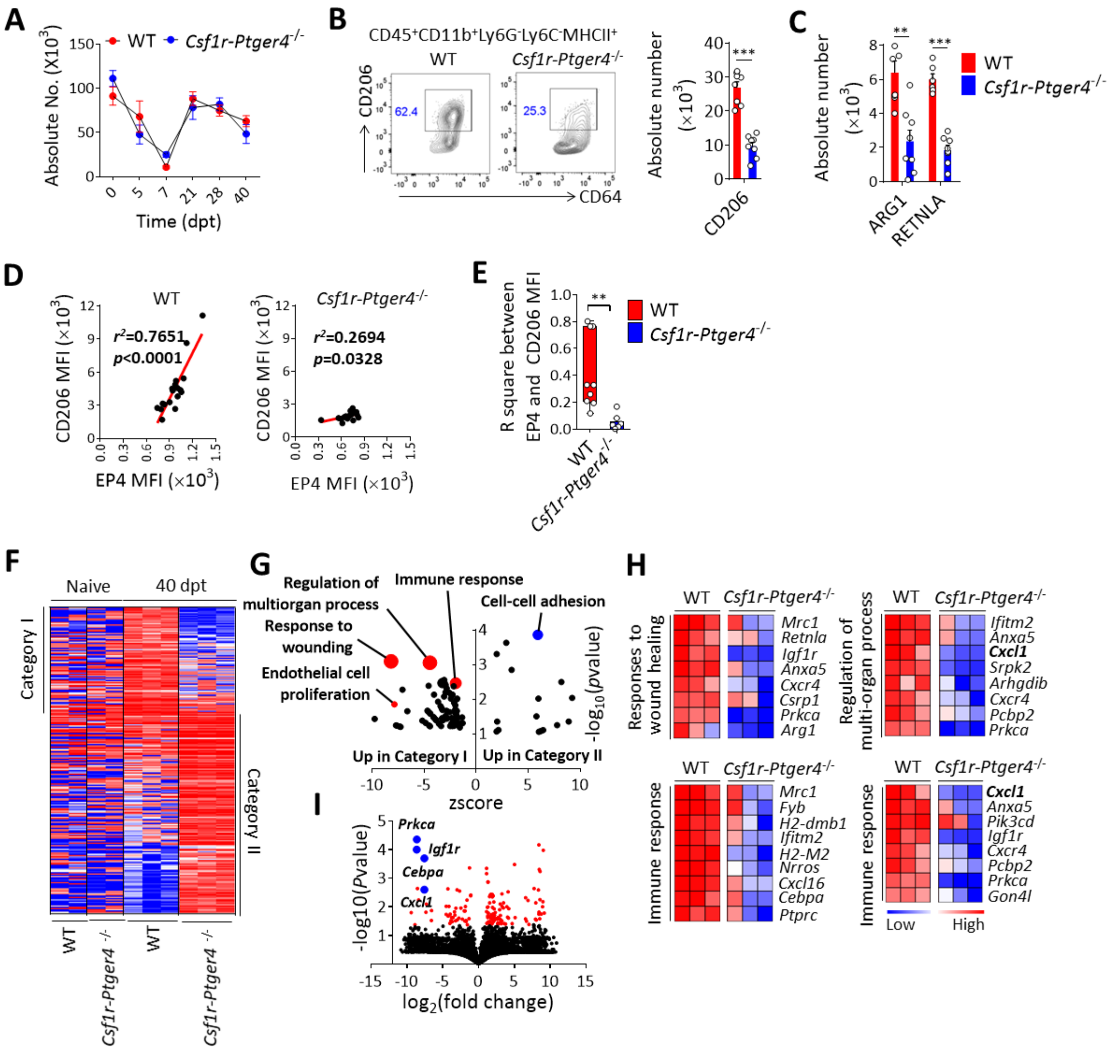


**Fig. 4. Wound healing phenotype of macrophages depends on the expression of EP4. A-H**, WT and *Csf1r-Ptger4*$^{-/-}$ mice were given 2.5 % DSS (w/v) for 5 d, followed by regular water, and analyzed at indicated time points. (**A**) Absolute number of mature colonic macrophages in WT and KO mice at indicated time points. (**B**) Representative contour plots of the CD206$^{+}$ macrophages (gated on the CD45$^{+}$CD11b$^{+}$Ly6G$^{-}$Ly6C$^{-}$MHCII$^{+}$ population) in the lamina propria of WT and KO mice at 40 dpt. Absolute number of CD206$^{+}$ macrophages (x10$^{6}$) of lamina propria cells is shown on the right. (**C**) Absolute number of ARG1$^{+}$ and RETNLA$^{+}$ macrophages in WT and

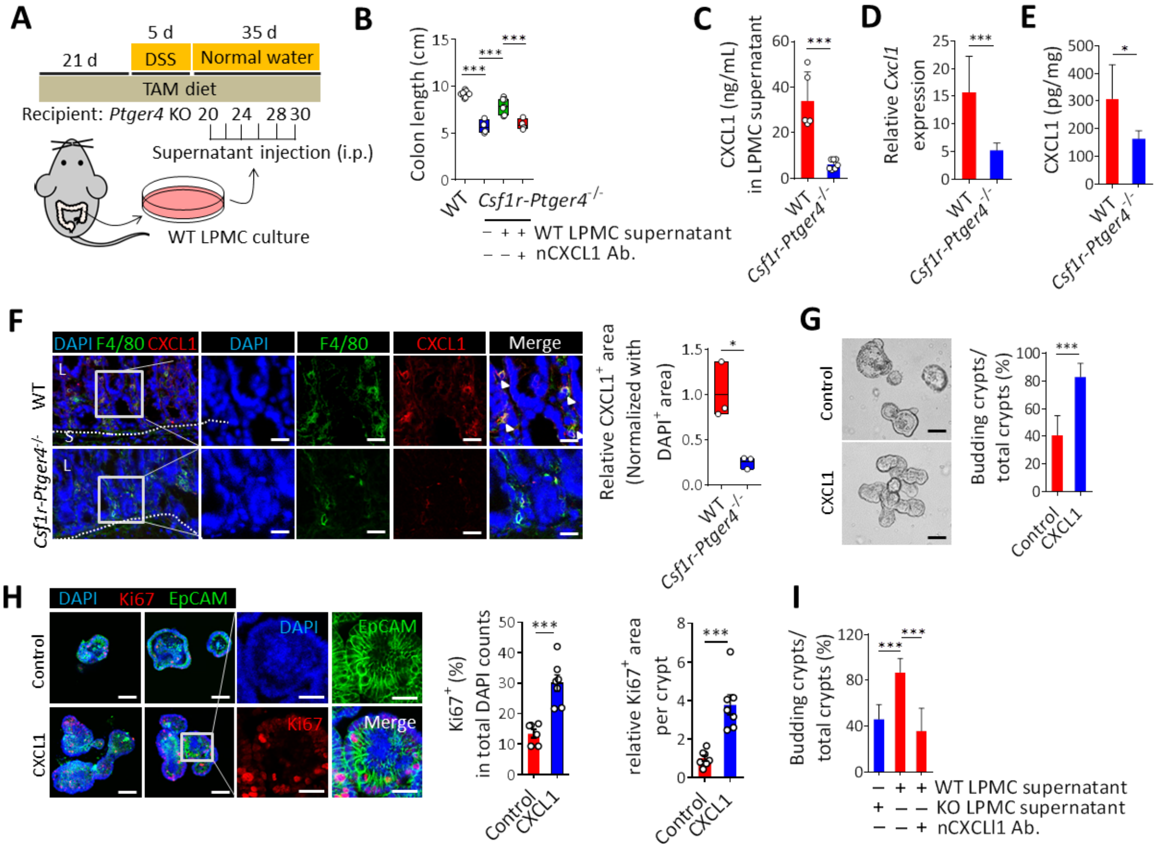


**Fig. 5. CXCL1 derived from EP4⁺ macrophages induces epithelial cell proliferation. A-E**, WT and *Csf1r-Ptger4*$^{-/-}$ mice were given 2.5 % DSS (w/v) for 5 d, followed by regular water, and analyzed at 40 dpt. Accordingly, 200 µL of culture supernatant of lamina propria mononuclear cells (LPMC) obtained from WT mice was intraperitoneally injected 6 times during the period of 20~30 dpt. Neutralizing antibodies for CXCL1 (20 µg/mL) was added to the supernatant before injection. (**A**) Overview of experimental set-up. (**B**) Comparison of colon lengths. (**C**) A total of 2 × $10^7$ LPMCs isolated from colons of WT and KO mice were cultured for 12 h. CXCL1 was determined in supernatant by enzyme-linked immunosorbent assay (ELISA). (**D**) mRNA expression of *Cxcl1* in whole colon lysates of WT and KO mice. (**E**) CXCL1 protein levels were determined in whole colon lysates of WT and KO mice. (**F**) Representative confocal images of WT and KO colons at 40 dpt

stained for F4/80 (green), CXCL1 (red), and DAPI (nuclei; blue) at 40 dpt.

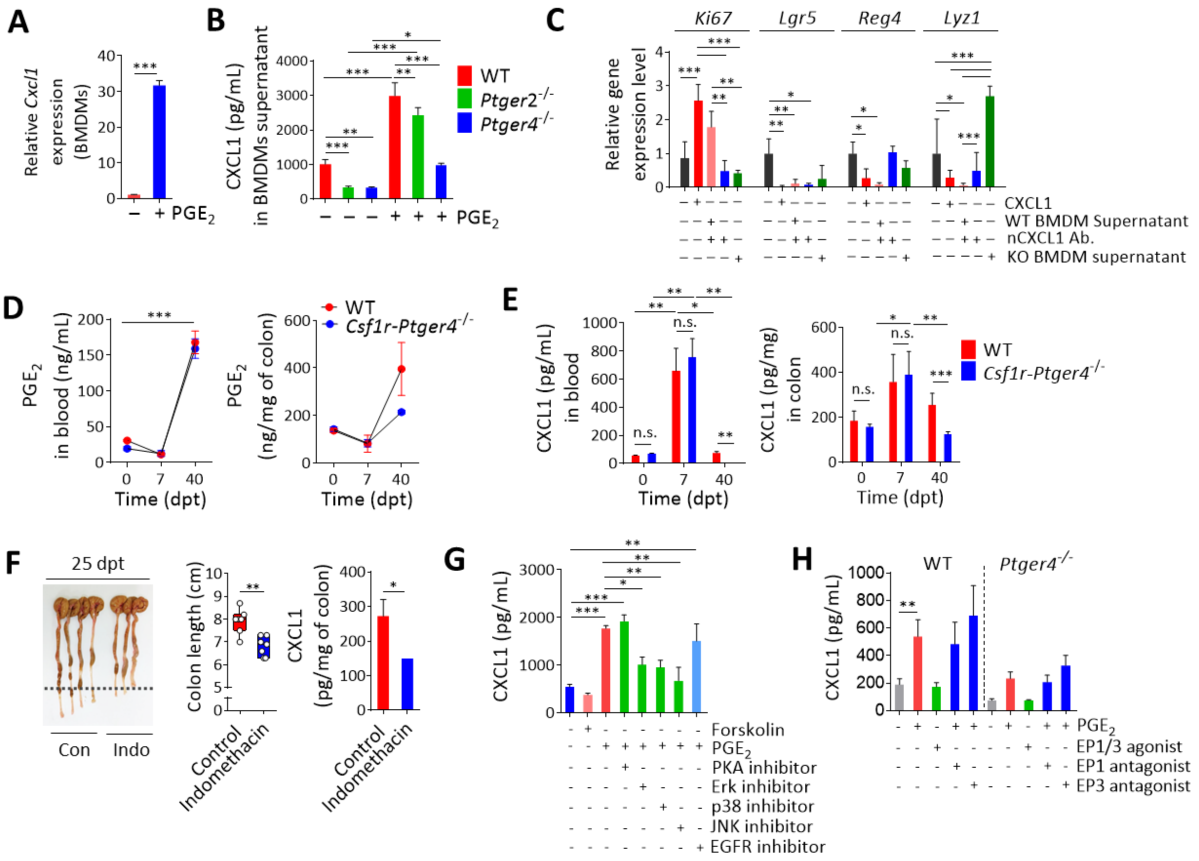


**Fig. 6. The $PGE_2$/EP4/MAPKs signaling pathway control secretion of Cxcl1 in macrophages.** (**A**) mRNA expression of *Cxcl1* in BMDMs treated with 10 µM of $PGE_2$. (**B**) A total of 5 × $10^5$ BMDMs obtained from WT, *Ptger2*$^{-/-}$ and *Ptger4*$^{-/-}$ mice were cultured for 3 d in the presence or absence of $PGE_2$. CXCL1 was determined in supernatant by ELISA. (**C**) Gene transcripts for *Ki67*, *Lgr5*, *Reg4*, and *Lyz1* of WT organoids were determined 2 d post treatment with 10 ng/mL CXCL1, WT BMDM sup, KO BMDM sup, 10 µg/mL of CXCL1 neutralizing antibody in WT BMDM supernatant. (**D**) The level of $PGE_2$ was measured in blood and whole colon lysates of WT and KO mice at different time points during DSS colitis using ELISA. (**E**) CXCL1 was determined in the blood and whole colon lysates of WT and KO mice at different time points during DSS colitis. (**F**) Comparison of colon lengths (left and middle) and CXCL1 in whole colon lysates (right) in WT mice treated with

indomethacin (10 mg/L in drinking water) from 15 until 25 dpt. (**G**) BMDMs were treated with forskolin (10 μM) or $PGE_2$ in the presence of inhibitors for PKA (H89, 20 μM), ERK (PD98059, 1 μM), p38 (SB203580, 0.1 μM), JNK (SP600125, 0.5 μM), or EGFR (erlotinib, 20 μM) and incubated for 3 d before measuring CXCL1 in the supernatant. (**H**) BMDMs were obtained from WT and KO mice and treated with $PGE_2$ in the presence of EP1 (SC-50189, 10 μM) or EP3 (L-798106, 10 μM) antagonist. The EP1/3 agonist, sulprostone, was added at a concentration of 10 μM. After 3 d, the level of CXCL1 in the supernatant was measured.

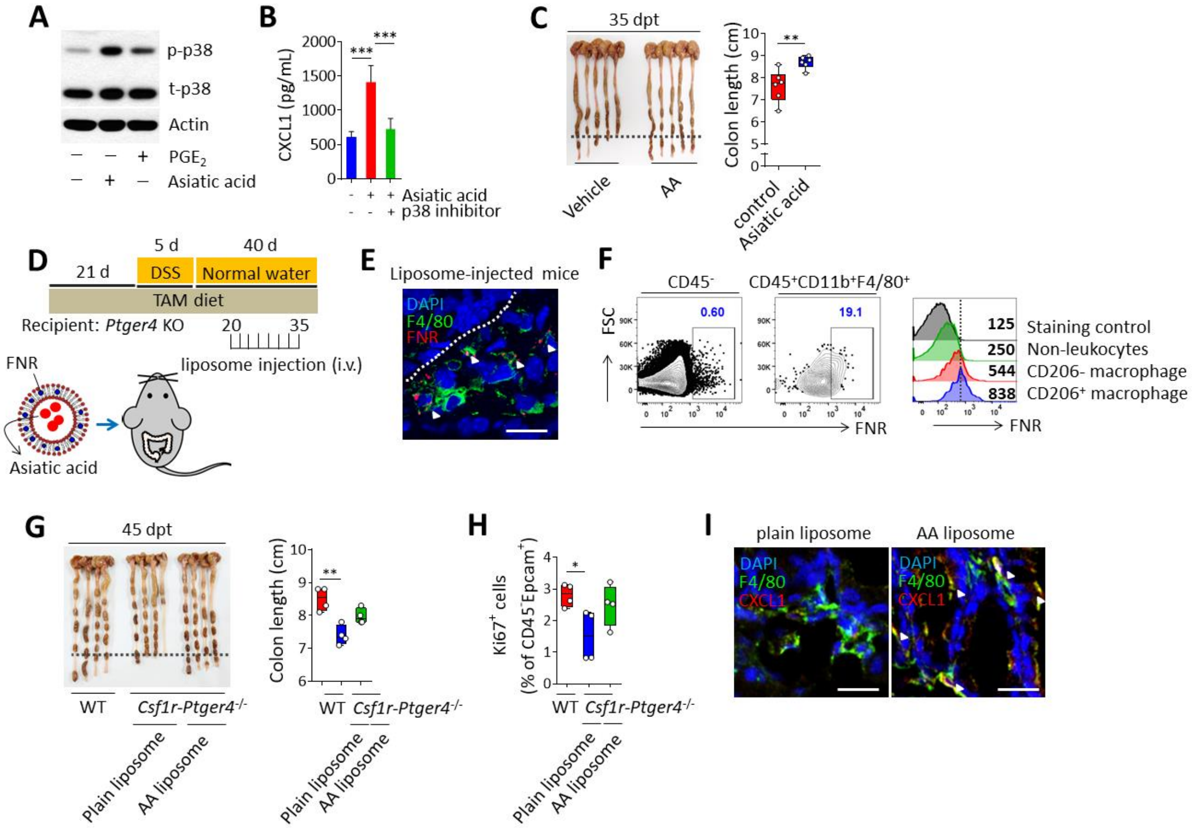


**Fig. 7. Therapeutic activation of MAPKs rescues intestinal regeneration of EP4-deficient mice.** (**A**) Expression of phospho-p38 determined by western blotting. BMDMs were treated with 20 µM of asiatic acid (AA) or 10 µM of $PGE_2$ for 1 h. (**B**) Determination of CXCL1 in supernatant was done by ELISA. BMDMs were treated with AA in the presence of a p38 inhibitor and cultivated for 3 d. (**C**) Comparison of colon lengths of WT mice (35 dpt) intraperitoneally injected with vehicle or AA (50 mg/kg) 8 times from 15 until 30 dpt during DSS-induced colitis. **D-I**, WT and *Csf1r-Ptger4*$^{-/-}$ mice were given 2.5 % DSS (w/v) for 5 d, followed by regular water, and analyzed at 45 dpt. FNR-conjugated, asiatic acid-containing liposomes were iv injected for a total of 8 times from 20 until 35 dpt. (**D**) Overview of experimental set-up. (**E**) Confocal image of colon of a WT mouse at 2 d post liposome injection stained for F4/80 (green), FNR (red), and DAPI (nuclei; blue)

**Funding:** S.S. was supported by the Basic Science Research Program through the National Research Foundation of Korea (NRF) funded by the Ministry of Science, ICT & Future Planning (2020R1A2C2010202). Y.N was supported by the NRF (NRF-2017R1D1A1B04031161). G.M. was supported by the FWO grant (G0D8317N, G0A7919N and S008419N), a grant from the International Organization for the Study of Inflammatory Bowel Diseases (IOIBD), a grant from the European Crohn´s and Colitis Organization (ECCO) and grants from the KU Leuven Internal Funds (C12/15/016 and C14/17/097). Collaboration between S.S. and G.M. laboratories is supported by a cooperation between FWO and NRF (VS03917N, VS07220N and 2019K2A9A1A06099778).

**Author contributions:** Y.N. and S.S. conceptualized this study; Y.N. and D.J. designed the experiments; Y.N., D.J., G.G., M.J., H.R., S.S., and J.P. performed the experiments; Y.N., D.J., H.H., B.P., S.L., H.L. and I.C. analyzed data; Y.N. and D.J. interpreted data; H.K., Y.K., T.S., S.R., K.P., J.I., and Y.L. provided vital reagents; Y.N., D.J., M.S., G.M., and S.S. contributed to manuscript writing; Y.N. wrote the manuscript.

**Competing interests:** The authors declare no competing interests.

**Data and materials availability:** Extended data will be available after publication.

**Correspondence and requests for materials** should be addressed to S.S. or G.M. Peer review information (please complete this sentence). Reprints and permissions information is available at (please complete this sentence).

**Supplementary Materials:** Materials and Methods and Figures S1-S8.

Supplementary Materials for

### E prostanoid receptor 4 expressing macrophages promote the regeneration of the intestinal epithelial barrier upon inflammation.

Yi Rang Na, Daun Jung, Michelle Stakenborg, Gyo Jeong Gu, Mi Reu Jeong, Hye Ri Jang, Soo Youn Suh, Hak Jae Kim, Yoon Hey Kwon, Tae Sik Sung, Seung Bum Ryoo, Kyu Joo Park, Jong Pil Im, Ji Yong Park, Yun Sang Lee, Heon Jong Han, Bo Youn Park, Sung Wook Lee, Ho Su Lee, Isabelle Cleynen, Gianluca Matteoli and Seung Hyeok Seok

**This PDF file includes:**

Materials and Methods

Figs. S1 to S8

Table S1 and S2.

#### Materials and Methods

Ethical statements

Human clinical samples were obtained from the Seoul National University Hospital after approval by the institutional review board (H-1104-066-358). Patients information is provided in Table S1.

Animal experiments were conducted in accordance with the Institute for Experimental Animal College of Medicine and performed according to the Guidelines for the Care and Use of Laboratory Animals by the Institutional Animal Care and Use Committee of Seoul National University (accession number SNU-171016-1-2).

Mice

Mice were housed and bred under specific pathogen free conditions at the animal facility of the Seoul National University Medical College. All mice were fed standard lab chow diet with access to water and food ad libitum. Male and female (10-12 wk old) Csf1r-cre/Esr1 (Tg(*Csf1r-Mer-iCre-Mer*)1Jwp strain) mice were purchased from Jackson Laboratory (Stock number 019098). B6.129S6(D2)-*Ptger4*$^{tm1.1Matb}$/BreyJ (EP4 floxed) mice were kindly provided by Shuh Narumiya (Kyoto University Graduate School of Medicine, Japan). B6.129-*Ptger2*$^{tm1Brey}$/JB6.(*Ptger2*$^{-/-}$) mice were purchased from Jackson Laboratory (Stock number 004376). SJL-*Ptprc*$^{a}$ *Pepc*$^{b}$/BoyJ mice (CD45.1 mice) were purchased from Jackson laboratory (Stock number 002014). B6.129S4-*Ccr2*$^{tm1Ifc}$/J (*Ccr2*$^{-/-}$ mice) were kindly provided by Prof. Jae-Hoon Choi in Hanyang University, Korea.

Csf1r-iCre mice were crossed with EP4 floxed mice to obtain Csf1r-Cre/Esr1 EP4$^{fl/fl}$ (*Csf1r-Ptger4*$^{-/-}$) mice. Littermate Cre-negative mice were used for matched wildtype (WT) in all experiments. To induce Cre-dependent deletion of EP4, tamoxifen containing diet (400 mg/kg, DooYeol Biotech, Korea) was administered for 21 d before starting of colitis and during the all experiments.

Dextran sodium sulfate-induced colitis model

Dextran sodium sulfate (DSS; molecular mass: 36 000-50 000 Da; MP Biomedicals, CA, USA) was added to drinking water at 2.5 % (w/v) for 5 d, followed by regular drinking water. Mice were monitored daily for weight loss, stool consistency, and hematochezia. Disease activity index (DAI) was set as the combined score of weight loss (score = 0; <1 %, 1; 1~5 %, 2; 5~10 %, 3: 10~20 %, 4: >20 %), stool blood (score = 0: absence, 2: presence, 4: gross bleeding), and stool consistency (score = 0: formed and hard, 1: formed but soft, 2: loose stools, 3: mild diarrhea, 4: gross diarrhea)(*1*). Mice were sacrificed in the $CO_2$ chamber at the indicated time points. In some experiments, anti-IL-4 (11B11, BioXcell, NH, USA) or rat IgG1k isotype antibodies (HRPN, BioXcell) were intraperitoneally injected 3 times a week throughout the whole disease recovery period (7~30 dpt) at a concentration of 250 μg per mice. Indomethacin was administrated in drinking water at 10 mg/L between 15~25 dpt.

Adoptive cell transfer

Bone marrow (BM) cells were obtained from WT mice and differentiated into bone marrow derived macrophages (BMDMs) for 7 d in RPMI 1640 media containing 10 % fetal bovine serum (FBS), 100 U/mL penicillin and 100 μg/mL streptomycin (Invitrogen, CA USA), and 2 mM of L-glutamine (Invitrogen), and supplemented with fresh recombinant murine colony-stimulating factor 1 (CSF1; 50 ng/mL; Miltenyi Biotec, CA, USA) every 3 d. For adoptive transfer of dendritic cells (DCs), bone marrow cells were cultured in complete medium with recombinant granulocyte-macrophage (GM)-CSF (20 ng/mL; Miltenyi Biotec) and interleukin 4 (IL-4; 5 ng/mL; Peprotech, NJ, USA). After 7 d of differentiation, cells were collected and $CD11c^+$ DCs were isolated using magnetic microbeads (Miltenyi Biotec). All cells were cultured at 37 °C in a humidified incubator containing 5 % $CO_2$. BMDMs were routinely >95 % $F4/80^+/CD11b^+$ cells, whereas DCs were routinely >90 % $CD11c^+$. Purified $2 \times 10^6$ cells were adoptively transferred into *Csf1r-Ptger4*$^{-/-}$ mice by intraperitoneal (IP) injection 3 times at 7, 10, and 13 d post treatment with DSS (dpt).

Bone marrow transplantation

For the generation of bone marrow chimeras, recipient ($CD45.1^+$) mice were irradiated with 6-MV x-rays from a linear accelerator (Varian Medical System, CA, USA) at a single dose of 12 Gy. The whole abdomen was shielded using a multileaf collimator. One day following irradiation, recipient mice were adoptively transferred with $1 \times 10^7$ CD45.2 bone marrow cells by lateral tail-vein injection derived from WT, *Ptger4*$^{-/-}$, *Ptger2*$^{-/-}$, and *Ccr2*$^{-/-}$ mice. After 7-8 weeks, DSS-colitis was induced in the bone marrow chimera mice with 2.5 % DSS in drinking water for 5 d.

Isolation of epithelial and lamina propria cells

After removing all excess fat and feces, colons were opened longitudinally, washed in Hank's balanced salt solution (HBSS; Invitrogen) supplemented with 1 % FBS, and cut into 0.5 cm sections. Tissue was incubated twice at 37 °C with shaking for 15 min in HBSS containing 1 mM EDTA and 1 % FBS. Strainer-filtrates were centrifuged (5 min, 300 *g*) to collect epithelial cells. Remaining tissue samples were minced and incubated with prewarmed 1× Minimum Essential Medium (MEM) α (Invitrogen) containing 2 mM L-glutamine, 100 μg/mL penicillin, 100 μg/mL streptomycin, 2-mercaptoethanol, and 10 % FBS, and supplemented with 1.25 mg/mL collagenase D (Roche, Basel, Swiss), 0.85 mg/mL collagenase V (Sigma), 1 mg dispase (Invitrogen), and 30 U/mL DNase (Roche Diagnostics GmbH) for 30–45 min in a shaking incubator at 37 °C. The resulting cell suspension was filtered through a 70 μm cell strainer (BD Falcon, NJ, USA) and lamina propria cells were collected after centrifugation (10 min, 300 *g*). To obtain lamina propria mononuclear cell (LPMC) culture supernatants, $2 \times 10^7$ of isolated single cells were cultured for 12 h in plain RPMI medium containing 100 U/mL penicillin and 100 μg/mL streptomycin (Invitrogen). Culture supernatants were collected, centrifuged (15 min, 500 *g*), and stored at -80 °C until used.

Quantitative real-time PCR

For quantitative real-time PCR analysis, total RNA was solubilized in TRIzol reagent (Invitrogen) and extracted according to the manufacturer's instructions. cDNA was synthesized from 1 mg of total RNA using reverse transcription, and the amount of mRNA was determined using real-time PCR analysis with the SYBR Green qPCR Pre Mix (Enzynomics, Daejeon, South Korea) on an ABI real-time PCR 7500 machine (Applied Biosystems, CA, USA). Samples were normalized to *Tbp* or *Rps18*. The primer sequences are described in Table S2.

Western Blotting

Western blotting was performed as previously described. Antibodies used were anti–EP4 (polyclonal, Cayman chemical, MI, USA) and anti-β-actin (C4, Santa-Cruz, TX, USA).

Flow cytometry

Single cell suspensions were stained with the following antibodies: EP4 (polyclonal) from Cayman chemical, F4/80 (BM8) and CD68 (KP1) from Santa-Cruz, CD45 (30-F11), CD11b (M1/70), Ly6G (1A8), IA/IE (M5/114.15.2), Ly6C (HK1.4), Arg1 (A1exF5), CD3 (145-2C11), CD115 (AFS98), and Epcam (G8.8) from Invitrogen, CD206 (C068C2), CD45.1 (A20), and CD45.2 (104) from Biolegend (CA, USA), CD64 (X54-5/7.1), and Retnla (RM0313-9A39) from NOVUS biologics (CO, USA), and cAMP (EP8471) from Abcam (Cambridge, UK), and Ki67 (D3B5) from Cell Signaling (Massachusetts, USA). Following incubation with purified anti-CD16/CD32 (93, Biolegend) for 10 min at 4 °C, cells were stained with appropriate antibodies at 4 °C in the dark. For intracellular staining, cells were fixed, permeabilized using 1 % paraformaldehyde (Merck, NJ, USA) and Perm/Wash buffer (BD Biosciences), and labeled with appropriate secondary antibodies. Data were acquired using the LSRFortessa system (BD Biosciences) and analyzed with the FlowJo software (Tree Star, OR, USA).

Myeloperoxidase activity measurement

Myeloperoxidase (MPO) activity was assessed to evaluate the extent of neutrophil infiltration into inflamed colonic lesion. Colon specimens were homogenized with 20 volume-equivalent of 50 mM phosphate buffer (pH 6.0) containing 0.5% hexadecyltrimethyl ammonium bromide (Sigma) using Precellys 24 (Bertin technologies) and sonicated for 10 s. Homogenates were freeze-thawed three times, and then centrifuged at 2,000 g for 15 min, 4 °C. Fourteen mL of supernatant was mixed with a solution of 1 mg/mL o-dianisidine hydrochloride (Sigma) and 0.0005% hydrogen peroxide. MPO activity was measured spectrophotometrically as the change in absorbance at 460 nm. One unit of MPO activity was defined as absorbance change per minute at 25 ℃ in the reaction with 1 mmol peroxidase.

Histology and confocal imaging

For histological analyses, tissues were flushed with PBS, then cut longitudinally and fixed in 10 % neutral buffered formalin, processed for wax embedding and routine hematoxylin and eosin staining. Images were acquired using a microscope (Eclipse Ci-L; NIKON) and features were blindly scored for the presence of mucosal erosion, ulceration, and leukocyte infiltration using the following grading system: Inflammation, 0 = no changes; 1= infiltrations in the lamina propria; 2 = extending into the submucosa; 3 = transmural extension; Epithelial damage, 0 = no change; 1 = loss of the basal one-third; 2 = loss of the basal two-third; 3 = entire crypt loss, 4 = epithelial erosion, 5 = confluent erosion; Mucosal architecture, 0 = no changes; 1 = 1 or 2 foci of ulcerations, 2 = 3 or 4 foci of ulcerations, 3 = confluent or extensive ulceration. For immunohistochemistry, slides were deparaffinized, rehydrated, and antigen-retrieved in a citrate buffer (10 mM sodium citrate, 0.05 % Tween 20, pH 6.0) for 15 min for further analysis. For detection of Ki67, tissues were incubated with anti-Ki67 (D3B5, Cell Signaling) for 12 h at 4 °C. Detection of primary antibody was achieved using the Dako Envision plus system.

For confocal imaging analysis, intestinal tissues were gently flushed with PBS and fixed with 4 % paraformaldehyde for 30 min. Tissues were embedded in optimal cutting temperature (OCT) compound, frozen and stored at -80 °C. Frozen tissues were sliced to 10 µm-thick sections and fixed in cold acetone for 10 min. Tissues were washed with PBS, then blocked with PBS containing 3 % bovine serum albumin (BSA) and 0.3 % Triton X-100 (Sigma-Aldrich) for 2 h at 25 °C. Tissue sections were then rinsed in staining buffer (1 % BSA and 0.3 % Triton X-100 in PBS) and incubated with anti-EP4 (polyclonal, Cayman chemicals), anti-CD68 (KP1; Santa-Cruz), anti-F4/80 (anti-MCA497GA; Bio-Rad), anti-Epcam (G8.8, Invitrogen), anti-Lgr5 (803420; R&D

systems, MN, USA), anti-Cxcl1 (48415; R&D systems), anti-Ki67 (D3B5, Cell Signaling), anti-CD206 (polyclonal, Abcam), and anti-Cxcr2 (polyclonal, Genetex, CA, USA). Tissues were stained overnight at 4 °C with antibodies diluted in PBS containing 1 % BSA and 0.3 % Triton X-100. Tissue sections were then washed 3 times, then stained with appropriate secondary antibodies and 4,6-diamidino-2-phenylindole (DAPI; Invitrogen) for 1 h. Tissue sections were washed 3 times and mounted with antifade mounting medium (Vectashield, CA, USA). Images were acquired on a Leica TCS SP8 confocal microscope (Wetzlar, Germany). For quantification of protein expressions, ImageJ was used to measure a positive signal from each fluorescently stained area.

Enzyme-linked immunosorbent assay (ELISA)

CXCL1 in cell culture supernatants, LPMC culture supernatants, blood or colon lysates were measured using the mouse Duoset ELISA kit (BD Biosciences) according to the manufacturer's protocol. To measure $PGE_2$, colons were homogenized in homogenization buffer (0.1 M phosphate, pH 7.4, containing 1 mM EDTA and 10 μM indomethacin) using a TissueLyser (Qiagen, Hilden, Germany) and centrifuged (15 min, 2,000 g) to obtain supernatants. $PGE_2$ was measured using ELISA Kits (Cayman chemicals).

Gene Expression Profiling with RNA-SeqIntestinal macrophages were purified by flow cytometric sorting (FACSAria III; BD Biosciences) at 0, 7, and 40 dpt from WT and *Csf1r-Ptger4*$^{-/-}$ mice. Respectively, 2 to 3 biological replicates were generated from the colon for each time point. RNA was isolated using TRIzol reagent (Invitrogen). Construction of the library was performed using the QuantSeq 3' mRNA-Seq Library Prep Kit (Lexogen, Vienna, Austria) according to the manufacturer's instructions. High-throughput sequencing was performed using single-end 75 sequencing on a NextSeq 500 system (Illumina, CA, USA). QuantSeq 3' mRNA-Seq reads were aligned using Bowtie2 (Langmead and Salzberg, 2012). Bowtie2 indices were either generated from genome assembly sequences or representative transcript sequences for aligning to the genome and transcriptome. The alignment file was used for assembling transcripts, estimating their abundances, and detecting the differential expression of genes. Differentially expressed genes were determined based on counts from unique and multiple alignments using coverage in Bedtools (Quinlan AR, 2010). The RT (Read Count) data were processed based on the global normalization method using the Genowiz™ version 4.0.5.6 (Ocimum Biosolutions, India). Gene classification and gene ontology analysis were based on searches done by DAVID (http://david.abcc.ncifcrf.gov/) and Medline databases (http://www.ncbi.nlm.nih.gov/). Enriched terms that passed FDR <20 % were reported.

Analysis of publicly available microarray and RNA-seq data

The transcriptome data were retrieved from Gene Expression Omnibus (GEO) database GSE36807 and GSE53306 (*2*). For microarray data, the series matrix files were downloaded, and the expression values were z-normalized for each sample. In the case of RNA-seq data, raw sequence read files were downloaded and filtered using fastp with default parameters (*3*). The remaining reads were then aligned to reference human transcriptome using kallisto to quantify expression levels (*4*). The reference human transcriptome annotation data (GRCh38.p13) were retrieved from GENCODE release 32 (*5*). Transcript-level expression values were merged into gene-level values by adding TPM values of all isoforms for each gene. Then, *Wilcoxon rank-sum test* was performed to compare expression levels of PTGER4 between patients with CD, UC and healthy controls.

Mouse intestinal organoids

Gut organoids were cultured according to a previously described protocol by Sato and Clevers (*6*). Briefly, crypts were harvested by incubating fragment of cut opened mouse small bowel in PBS containing 2 mM EDTA. The epithelium was released by vigorous shaking and crypts were separated using a 70 µm strainer. Crypts were seeded in growth factor reduced Matrigel (BD Biosciences) and grown in Intesticult medium (Stem cell Technologies, Canada) at 37 °C in a 5 % $CO_2$ atmosphere. Next day, LPMC culture supernatants or 10 ng/mL recombinant CXCL1 (Peprotech) was added to the culture medium. Fresh medium was replaced every 4 d of culture. At 7 d of cultivation, organoids were photographed, and quantitative statistics of the number of budding crypts were obtained. To examine gene transcripts, organoids were treated with BMDM sup or recombinant CXCL1 at 5 d of culture and harvested for further analysis 2 d post treatment. To observe phosphorylation of CXCR2, organoids were treated with recombinant CXCL1 and analysed after 2 d of treatment.

Bone marrow-derived macrophages

Bone marrow cells were obtained from the femur and tibia of *Csf1r-Ptger4*$^{-/-}$ and ittermate control mice, or B6.129-*Ptger2*$^{tm1Brey}$/JB6.(*Ptger2*$^{-/-}$) mice and differentiated into bone marrow derived macrophages (BMDMs) for 7 d in RPMI 1640 media containing 10 % fetal bovine serum (FBS), 100 U/mL penicillin and 100 µg/mL streptomycin (Invitrogen, CA USA), and 2 mM of L-glutamine (Invitrogen), and supplemented with fresh recombinant murine colony-stimulating factor 1 (CSF1; 50 ng/mL; Miltenyi Biotec, CA, USA) every 3 d. Cells were treated with 4-hydroxytamoxifen (6 µM ; Sigma) for the last 3 d of differentiation, and media replaced with fresh CSF1-containing culure media before further chemical treatments. In same experiment BMDMs were pretreated with 100 μM indomethacin 1 h before $PGE_2$ stimulation.

Chemical activators and inhibitors

Inhibitors for protein kinase A (H89), ERK (PD98059), p38 (SB203580), JNK (SP600125), and EGFR (erlotinib) were purchased from Cayman chemical (Ann Arbor, USA). An activator for adenylyl cyclase (forskolin) was obtained from Cayman chemical. An activator for p38 (Asiatic acid) was obtained from Sigma (St. Louis, USA). Agonists for EP1/3 (sulprostone), EP1 (SC-50189), and EP3 (L-798106) were obtained from Cayman chemical.

Liposomes

The hydrodynamic diameter and size distribution of nanoparticles were analyzed by a dynamic light scattering (DLS) system Zetasizer Nano ZS90 (Malvern Instruments Ltd, Worcestershire, UK). Clickable liposomes (DBCO-Liposome) were prepared by mixing a predetermined ratio of distearoylphosphatidylcholine (DSPC, Avanti Polar Lipid), cholesterol (Avanti Polar Lipid), and distearoylphosphatidylethanolamine (DSPE, Avanti Polar Lipid) with 2.5 mol% DBCO-PEG2000-DSPE (Avanti Polar Lipid), and then the crude mixture was extruded. To conjugate Mannose and FNR on the liposomal surface, 6 μM of Mannose-azide (α-MAN-TEG-$N_3$, Iris Biotech GmbH, Marktredwitz, Germany) and FNR-azide were added to 10 mL of DBCO-liposome solution (10 mg/mL), followed by a purification with PD-10 desalting column (Mannose-Liposome). To make liposomes with AA inserted in the membrane, lipids were mixed and dried with AA dissolved in ethanol (Mannose-Liposome with AA).

Statistics

All data are presented as average values with standard error of the mean (SEM). Mann-Whitney (two-tailed) *U*-test, One-way ANOVA, and Two-way ANOVA test were used to determine statistical significance. Calculations and graphs were performed using the GraphPad Prism 6 software (CA, USA). All experiments were performed at least twice, with similar results obtained each time. Figure legends denote the specific statistical tests used for each experiment. Statistical significance was determined as $P<0.05$.

Data availability

RNA seq data that support the findings of this study have been deposited in the Gene Expression Omnibus repository with the accession code GSE141093.

**Table S1.** Characteristics of the human sample used in this study

| Case No. | Diagnosis | Enrolled date | Age | Sex | Clinical activity score[a] | Biopsy site[b] | MRC1⁺ % co-localized with CD68 | EP4⁺ % co-localized with CD68 | Sample type |
|---|---|---|---|---|---|---|---|---|---|
| 1 | Normal (colon polyp) | 10-21-13 | 60 | M | | A-colon | 86.9565 | 85.7143 | Paraffin block |
| 2 | Normal (colon lipoma) | 10-29-13 | 60 | F | | A-colon | 44.4444 | 45 | Paraffin block |
| 3 | Normal (colon polyp) | 11-26-13 | 56 | M | | A-colon | 88.8889 | 86.6667 | Paraffin block |
| 4 | Normal (CD, least likely) | 12-12-13 | 21 | M | | A-colon | 72 | 80 | Paraffin block |
| 5 | Active UC | 11-30-16 | 49 | M | Partial 5+2 | A-colon | 48 | 84.6154 | Paraffin block |
| 6 | Active UC | 01-24-18 | 22 | F | Partial 3+2 | A-colon | 25.5814 | 8.51064 | Paraffin block |
| 7 | Active UC | 03-29-17 | 22 | M | Partial 5+0 | T-colon | 21.875 | 15.625 | Paraffin block |
| 8 | Active UC | 10-23-17 | 26 | M | Partial 7+2 | A-colon | 14.2857 | 19.0476 | Paraffin block |
| 9 | Active CD | 10-23-17 | 30 | M | 151 | A-colon | 25 | 28.5714 | Paraffin block |
| 10 | Active CD | 06-25-18 | 28 | M | 413.74 | T-colon | 12.8205 | 29.6296 | Paraffin block |
| 11 | Active CD | 09-28-17 | 20 | M | 172 | Terminal ileum | 35.2941 | 48.3871 | Paraffin block |
| 12 | Active CD | 01-03-18 | 47 | M | 153 | A-colon | 26.6667 | 31.5789 | Paraffin block |
| 13 | Active CD | 09-05-16 | 16 | F | 393.75 | A-colon | 7.69231 | 25.641 | Paraffin block |
| 14 | Active CD | 02-21-17 | 25 | F | 172 | Terminal ileum | 33.3333 | 15 | Paraffin block |
| 15 | UC in remission | 04-11-18 | 25 | M | Partial 0+0 | T-colon | 28.9474 | 60 | Paraffin block |
| 16 | UC in remission | 03-21-18 | 47 | F | Partial 0+0 | T-colon | 65.5172 | 100 | Paraffin block |
| 17 | UC in remission | 12-27-17 | 47 | M | Partial 0+0 | T-colon | 73.1707 | 62.5 | Paraffin block |
| 18 | UC in | 07-31-17 | 46 | M | 0 | T-colon | 68.5714 | 60 | Paraffin |

| | | | | | | | | | |
|---|---|---|---|---|---|---|---|---|---|
| | remission | | | | | | | | block |
| 19 | CD in remission | 07-31-17 | 32 | M | 19.28 | Terminal ileum | 69.2308 | 83.3333 | Paraffin block |
| 20 | CD in remission | 10-01-15 | 56 | M | 115 | Terminal ileum | 57.8947 | 94.1176 | Paraffin block |
| 21 | CD in remission | 07-11-17 | 39 | F | 42.72 | A-colon | 80 | 83.3333 | Paraffin block |
| 22 | CD in remission | 06-28-18 | 42 | F | 55.88 | T-colon | 91.6667 | 90 | Paraffin block |
| 23 | CD in remission | 07-12-16 | 35 | M | 44.4 | T-colon | 95.7447 | 63.8298 | Paraffin block |
| 24 | CD in remission | 12-28-15 | 16 | M | 129 | A-colon | 66.6667 | 100 | Paraffin block |
| 25 | Normal (colon cancer) | 02-20-20 | 65 | M | | A-colon | Flow cytometry | Flow cytometry | Fresh tissue |
| 26 | Normal (rectal cancer) | 02-20-20 | 67 | F | | S-colon | Flow cytometry | Flow cytometry | Fresh tissue |
| 27 | Normal (rectal cancer) | 02-24-20 | 60 | M | | S-colon | Flow cytometry | Flow cytometry | Fresh tissue |
| 28 | Normal (rectal cancer) | 02-25-20 | 66 | M | | S-colon | Flow cytometry | Flow cytometry | Fresh tissue |
| 29 | Active UC | 12-24-19 | 31 | M | | S-colon | Flow cytometry | Flow cytometry | Fresh tissue |
| 30 | Active CD | 04-01-20 | 42 | F | 375 | A-colon | Flow cytometry | Flow cytometry | Fresh tissue |
| 31 | Active CD | 04-20-20 | 33 | M | 323 | A-colon | Flow cytometry | Flow cytometry | Fresh tissue |

[a]Score was assessed according to the mayo scoring system in case of ulcerative colitis (UC)(*1*) or the Crohn's disease activity index (CDAI)(*2*) in case of CD.
[b]A-colon; ascending colon, T-colon; transverse colon, S-colon; sigmoid colon

**Table S2.** Primer sequences used for gene amplification in this study

| Gene (for qRT-PCR) | Primers |
|---|---|
| *Ptger4* (exon2) | Forward: TGA CCC AAG CAG ACA CCA CCT<br>Reverse: TCC CAC TAA CCT CAT CCA CCA A |
| *Lgr5* | Forward: GGG AGC GTT CAC GGG CCT TC<br>Reverse: GGT TGG CAT CTA GGC GCA GGG |
| *Bmi1* | Forward: AAT TAG TTC CAG GGC TTT TCA A<br>Reverse: CTT CAT CTG CAA CCT CTC CTC TAT |
| *Muc2* | Forward: CAG TTT ATT CCT GTG TGC CCA AGG<br>Reverse: GGC TTC AGA ATA ATG TAC TGC TGC |
| *Reg4* | Forward: CTG GAA TCC CAG GAC AAA GAG TG<br>Reverse: CTG GAG GCC TCC TCA ATG TTT GC |
| *Lyz1* | Forward: GAG ACC GAA GCA CCG ACT ATG<br>Reverse: CGG TTT TGA CAT TGT GTT CGC |
| *Lyz2* | Forward: ATG GAA TGG CTG GCT ACT ATG G<br>Reverse: ACC AGT ATC GGC TAT TGA TCT GA |
| *Cxcl1* | Forward: TCC AGA GCT TGA AGG TGT TGC C<br>Reverse: AAC CAA GGG AGC TTC AGG GTC A |
| *Arg1* | Forward: GTC TGG CAG TTG GAA GCA TCT<br>Reverse: GCA TCC ACC CAA ATG ACA CA<br>Probe: TGG CCA CGC CAG GGT CCA C |
| *Mrc1* | Forward: AAT GAA GAT CAC AAG CGC TGC<br>Reverse: TGA CAC CAG CGG AAT TTC T |
| *Retnla* | Forward: CCT GCT GGG ATG ACT GCT ACT<br>Reverse: TCC ACT CTG GAT CTC CCA AGA<br>Probe: TGT GCT TGT GGC TTT GCC T |
| *Rps18* | Forward: GCA ATT ATT CCC CAT GAA CG<br>Reverse: GGC CTC ACT AAA CCA TCC AA |
| *Tbp* | Forward: AAG GGA GAA TCA TGG ACC AG<br>Reverse: CCG TAA GGC ATC ATT GGA CT |
| **Gene (for genotyping)** | **Primers** |
| *Csf1r* transgene | Forward: AGA TGC CAG GAC ATC AGG AAC CTG<br>Reverse: ATC AGC CAC ACC AGA CAC AGA GAT C- |
| Internal control | Forward: CTA GGC CAC AGA ATT GAA AGA TCT<br>Reverse: GTA GGT GGA AAT TCT AGC ATC ATC C |
| EP4 floxed | Forward: TCT GTG AAG CGA GTC CTT AGG CT<br>Reverse: GTT AGA TGG GGG GAG GGG ACA ACT |

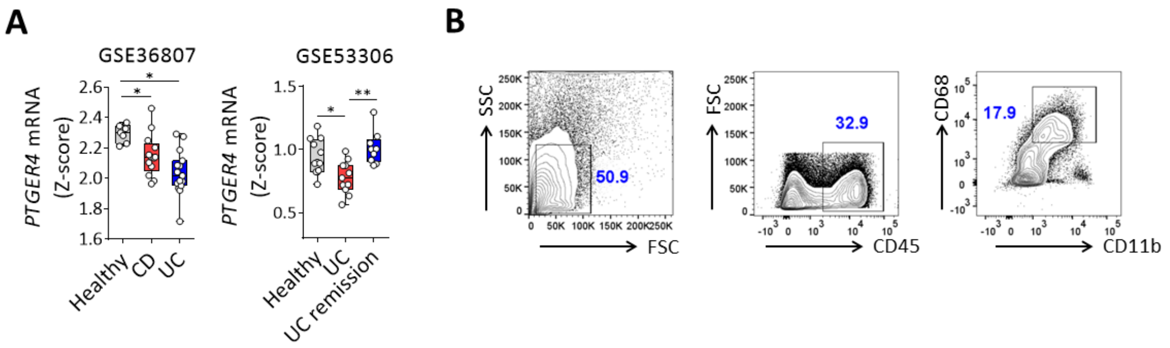


**Fig. S1. Analysis of the expression of EP4 in human patients with IBD.** (**A**) *PTGER4* mRNA levels in healthy control, UC and CD patients from the GSE36807 and GSE53306. Box and whisker plots indicate the median and the 25th and 75th percentiles, with minimum and maximum values at the extremes of the whiskers. CD, Crohn's disease, UC, ulcerative colitis. One-way ANOVA was performed. *$P$ < 0.05; **$P$ < 0.01. (**B**) Representative contour plots for the gating strategy of human colonic mucosal macrophages. $CD45^+CD11b^+CD68^+$ cells were further analyzed for the expression of EP4 and CD206. n = 3 of individual cases.

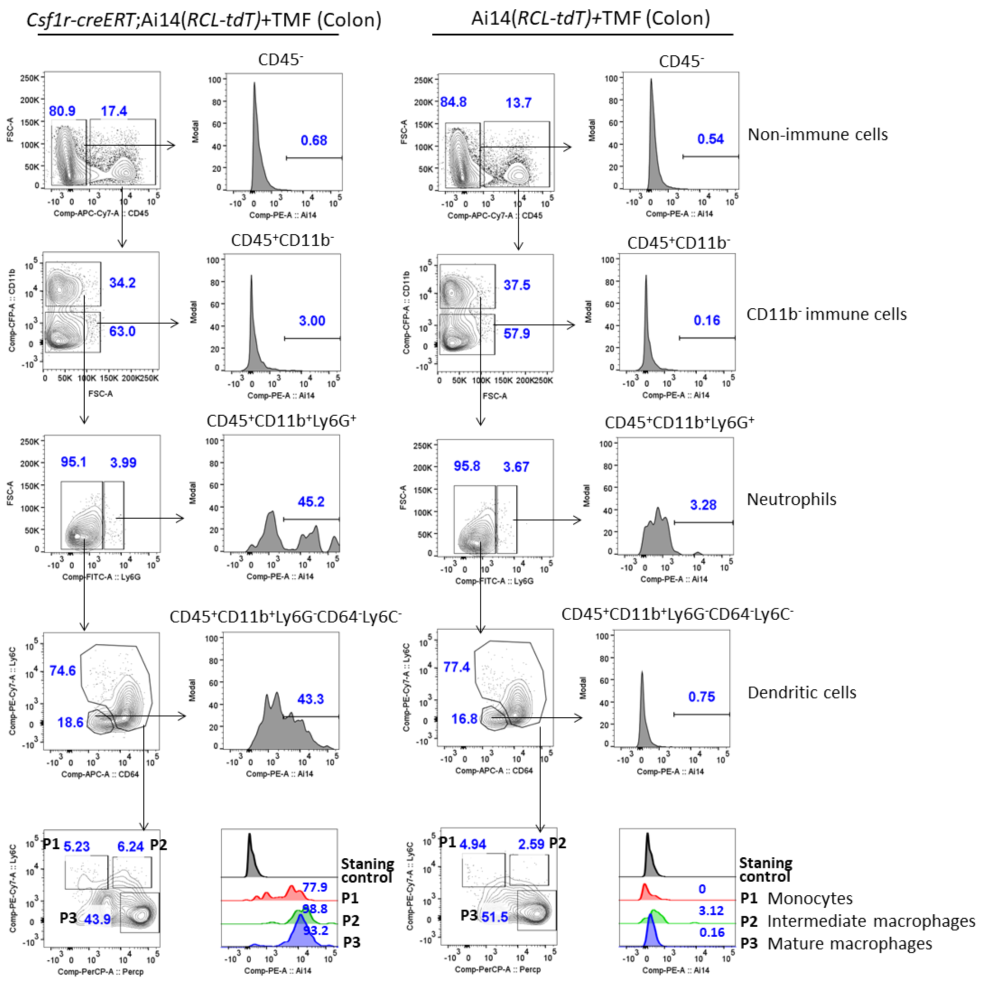


**Fig. S2. Expression of CSF1R in major cell populations of the large intestine.** Tamoxifen (TMF, 5 mg/mL in corn oil) was orally injected into *Csf1r-creERT*;Ai14(*RCL-tdT*) and Ai14(*RCL-tdT*) mice for 5 consecutive days and then mice were sacrificed. Colonic lamina propria cells were isolated and analyzed by flow cytometry. Gating was performed according to the following gating strategy: $CD45^-$ (non-immune cells), $CD45^+CD11b^-$ ($CD11b^-$ immune cells), $CD45^+CD11b^+Ly6G^+$ (neutrophils), $CD45^+CD11b^+Ly6G^-Ly6C^-CD64^-$ (dendritic cells), $CD45^+CD11b^+Ly6G^-Ly6C^+MHCII^-$ (P1; monocytes), $CD45^+CD11b^+Ly6G^-Ly6C^+MHCII^+$ (P2; intermediaries), and $CD45^+CD11b^+Ly6G^-Ly6C^-MHCII^+$ (P3; mature macrophages). mCherry signals were detected and shown in histograms.

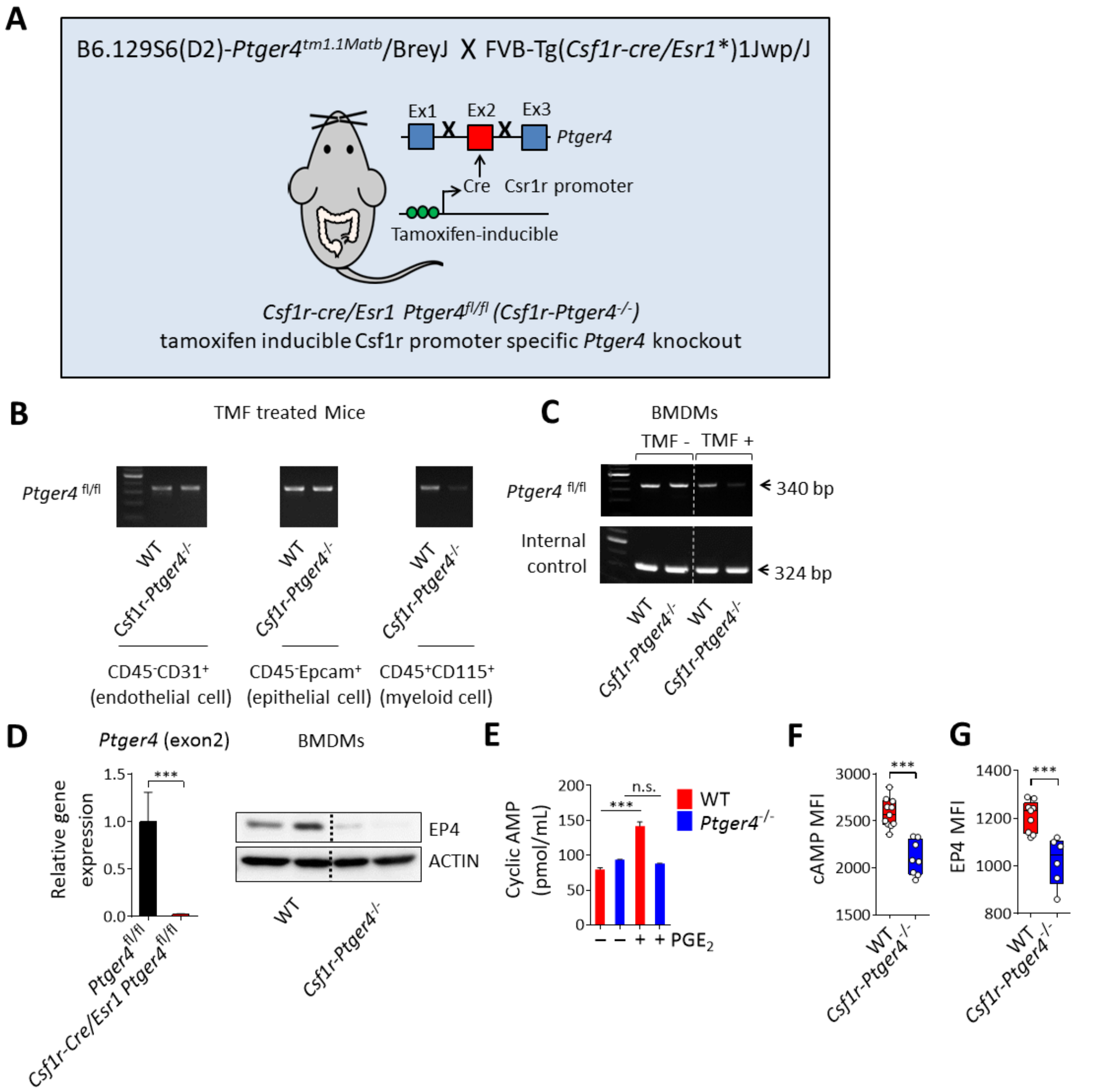


**Fig. S3. Generation and confirmation of specific knock out in *Csf1r-Ptger4*$^{-/-}$ mice.** (**A**) Schematic representation of genetic construct in *Csf1r-Ptger4*$^{-/-}$ mice. Deletion of EP4 was achieved by crossing Csf1r-cre/Esr1 mice with EP4 floxed mice. (**B**) Analysis of gene recombination in the DNA isolated from sorted CD45$^{-}$CD31$^{+}$ endothelial cells, CD45$^{-}$EpCAM$^{+}$ epithelial cells, and CD45$^{+}$CD115$^{+}$ myeloid cells obtained from tamoxifen injected *Csf1r-Ptger4*$^{-/-}$ mice. PCR products for the floxed EP4 allele (340 bp) were detected using gel electrophoresis.

(**C**) BMDMs were obtained from WT and KO mice and differentiated with or without supplementation with 4-hydroxytamoxifen (TMF, 6 μM). Amplification of deleted sequences was examined using PCR. (**D**) Real time quantitative PCR to determine the mRNA of exon2 of EP4 in BMDMs obtained from WT and *Csf1r-Ptger4*$^{-/-}$ mice (left). EP4 protein was detected by western blotting in 4-hydroxytamoxifen-treated BMDMs (right). (**E**) cAMP was determined in cell lysates using ELISA. BMDMs were obtained from WT and *Ptger4*$^{-/-}$ mice and stimulated with 10 μM of $PGE_2$ for 20 min. (**F**) Intracellular cAMP was detected by flow cytometry in colonic mature macrophages of WT and KO mice. (**G**) The mean fluorescent intensity of EP4 was measured by flow cytometry in colonic macrophages of WT and KO mice. MFI, mean fluorescence intensity. In panels D, F, and G, statistical significance was determined by a two-tailed unpaired Student's *t*-test. In panel E, one-way ANOVA was performed. *$P < 0.05$; ***$P < 0.001$.

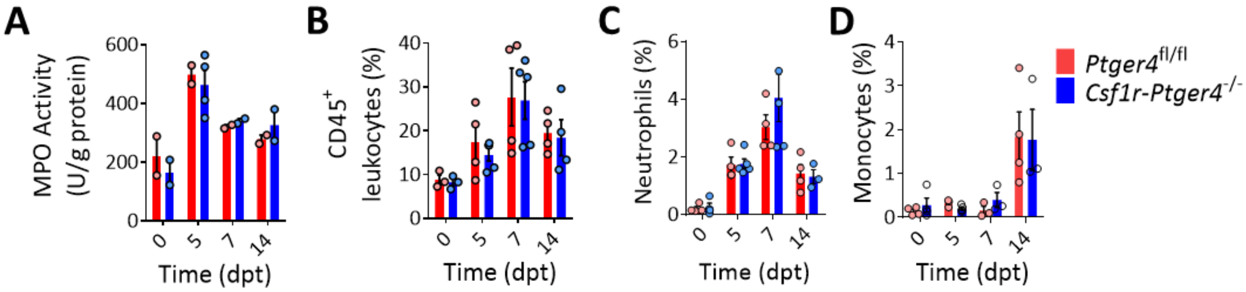


**Fig. S4. Macrophage specific deficiency of EP4 does not contribute to the extent of inflammation in a DSS-induced colitis.** (**A**) Myeloperoxidase (MPO) activity was measured in the colon lysates of WT and *Csf1r-Ptger4*$^{-/-}$ mice at indicated time points. **B-D**, The frequency of total leucocytes (**B**), neutrophils (**C**), and monocytes (**D**) in colonic lamina propria cells isolated from WT and *Csf1r-Ptger4*$^{-/-}$ mice at different time points during DSS colitis. Statistical significance was determined by two-way ANOVA.

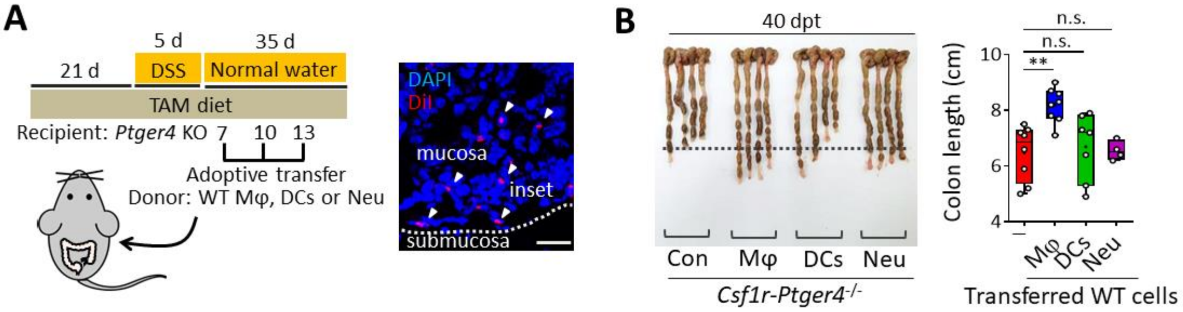


**Fig. S5. EP4⁺ macrophages contribute to complete resolution of colitis. A-B**, Adoptive transfer of $2 \times 10^6$ WT macrophages (Mφ), dendritic cells (DCs) and neutrophils (Neu) was performed in *Csf1r-Ptger4*$^{-/-}$ mice in a total of 3 times during the resolving phase of colitis. (**A**) Overview of experimental set-up is depicted on the left. Confocal image shows DiI-labeled Mφ (arrowheads) in the colon of recipient mice 7 d post injection. Scale bar, 20 µm. (**B**) Comparison of colon lengths of KO mice injected with Mφ, DCs or Neu at 40 dpt. One-way ANOVA was performed. $^{**}P < 0.01$.

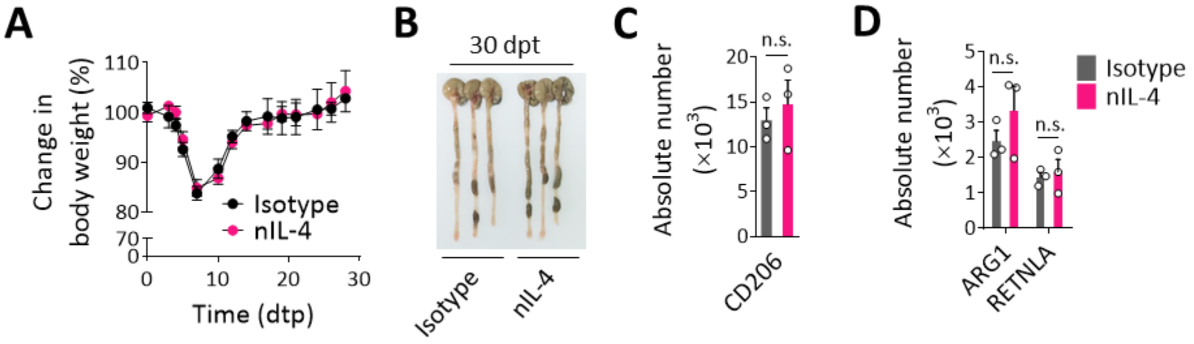


**Fig. S6. Wound healing phenotype of intestinal macrophages does not depend on the IL-4 during DSS colitis. A-D**, WT mice were given 2.5 % DSS (w/v) for 5 d, followed by regular water, and analyzed at indicated time points. 250 µg of IL-4 neutralizing antibody or corresponding isotype antibody was intraperitoneally injected 3 times a week throughout the whole recovery period of DSS colitis. (**A**) Body weights are shown as a percentage of the initial weight. n = 5 mice. (**B**) Comparison of colon lengths at 30 dpt. **C-D,** Absolute number of $CD206^+$ (**C**), $ARG1^+$, and $RETNLA^+$ macrophages (**D**) in a total of $10^6$ lamina propria isolates analyzed by flow cytometry. A two-tailed unpaired Student's *t*-test was performed. n.s., not significant.

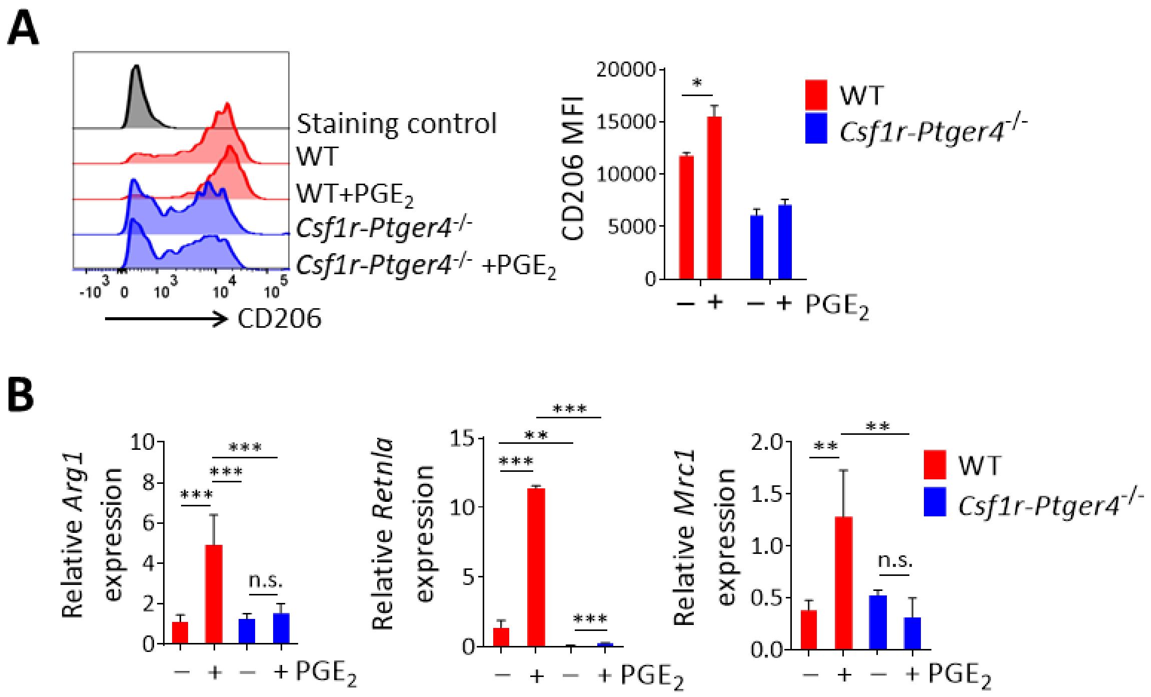


**Fig. S7. $PGE_2$ induces M2 macrophage markers in an EP4-dependent manner.** (**A**) Histograms showing the expression of CD206 in BMDMs analyzed by flow cytometry (left). BMDMs obtained from WT and *Csf1r-Ptger4*$^{-/-}$ mice were treated with 10 μM of $PGE_2$ for 2 d. Quantitative graph for CD206 is shown in right. MFI, mean fluorescence intensity. (**B**) BMDMs obtained from WT and *Csf1r-Ptger4*$^{-/-}$ mice were treated with 10 μM of $PGE_2$ for 6 h. Expression levels of *Arg1*, *Retnla*, and *Mrc1* were determined by real-time PCR. Statistical significance was determined by one-way ANOVA. *$P < 0.05$; **$P < 0.01$; ***$P < 0.001$.

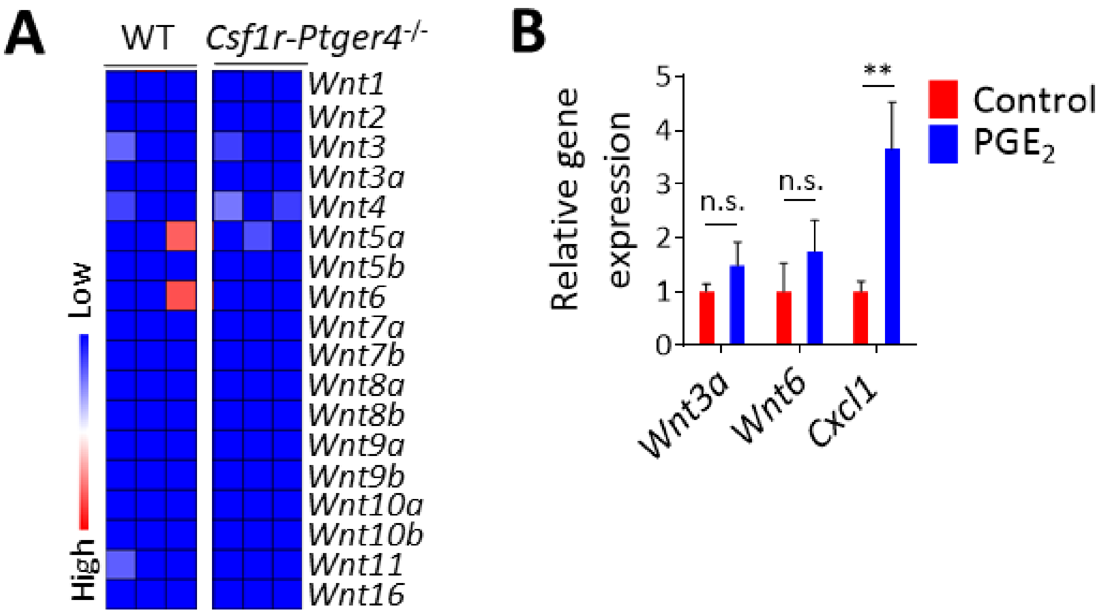


**Fig. S8. PGE$_2$/EP4 signaling is not involved in Wnt synthesis in mature macrophages.** (**A**) WT and *Csf1r-Ptger4*$^{-/-}$ mice were given 2.5 % DSS (w/v) for 5 d, followed by regular water, and analyzed. Heatmap depicts the expressions of *Wnt* genes between WT and KO at 40 dpt. (**B**) BMDMs obtained from WT mice were treated with 10 µM of PGE$_2$ for 8 h. Expression levels of *Wnt3a*, *Wnt6*, and *Cxcl1* were determined by real-time PCR. Statistical significance was determined by one-way ANOVA. $^{**}P < 0.01$. n.s; not significant.

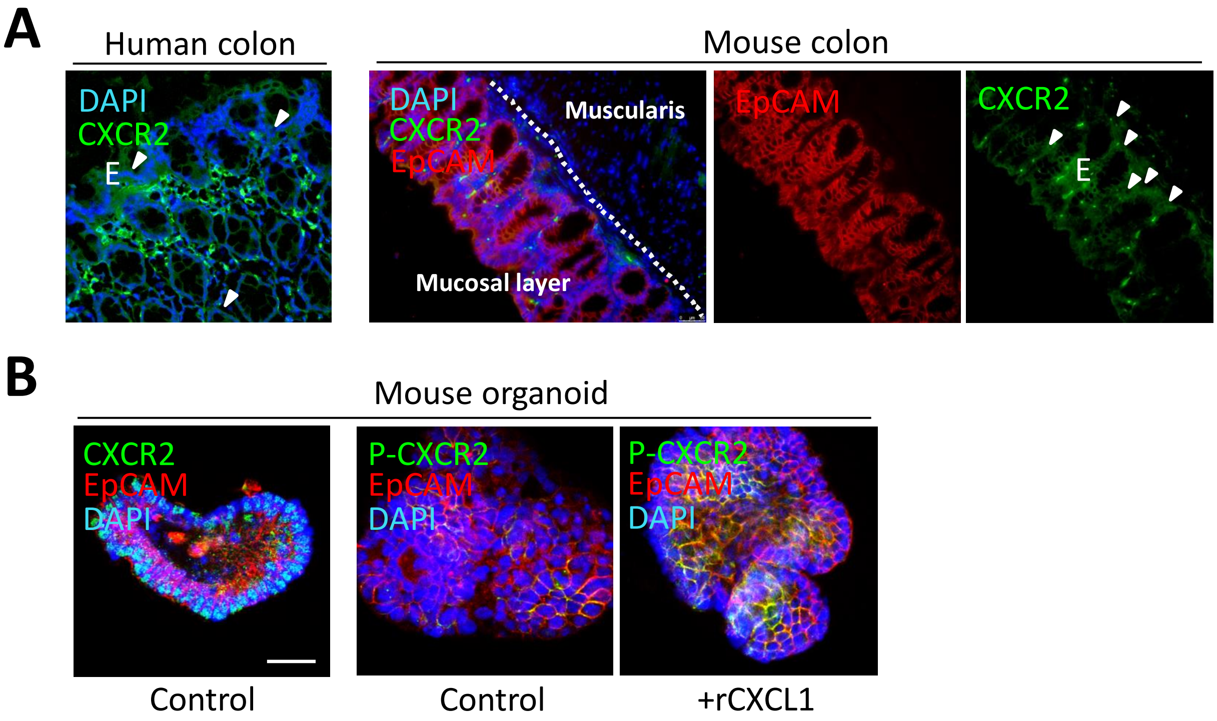


**Fig. S9. Human and mouse colonic epithelial cells express CXCR2.** (**A**) Representative confocal images of the human and mouse colon stained for CXCR2 (green), EpCAM (red, in case of mice), and DAPI (nuclei; blue). Arrowheads indicate $CXCR2^{+}$ epithelial cells. (**B**) Representative confocal images of organoids stained for CXCR2 (green) or phospho-CXCR2 (green) with parallel staining for EpCAM (red) and DAPI (nuclei; blue). Organoids were treated with CXCL1 for 1 h.
